# Warning coloration, body size and the evolution of gregarious behavior in butterfly larvae

**DOI:** 10.1101/2022.07.22.500975

**Authors:** Callum F McLellan, Stephen H Montgomery, Innes C Cuthill

**Affiliations:** University of Bristol

**Keywords:** comparative analysis, phylogenetics, gregarious behavior, aposematism, body size, defensive coloration

## Abstract

Many species gain anti-predator benefits by combining gregarious behavior with warning coloration, yet there is debate over which trait evolves first, and which is the secondary adaptive enhancement. Body size can also influence how predators receive aposematic signals, and potentially constrain the evolution of gregarious behavior. To our knowledge, the causative links between the evolution of gregariousness, aposematism and larger body sizes have not been fully resolved. Here, using the most recently resolved butterfly phylogeny and an extensive new dataset of larval traits, we reveal the evolutionary interactions between important traits linked to larval gregariousness. We show that larval gregariousness has arisen many times across the butterflies, and aposematism is a likely prerequisite for gregariousness to evolve. We also find that body size may be an important factor for determining the coloration of solitary, but not gregarious larvae. Additionally, by exposing artificial ‘larvae’ to wild avian predation, we show that undefended, cryptic ‘larvae’ are heavily predated when aggregated but benefit from solitariness, whereas the reverse is true for aposematic prey. Our data reinforce the importance of aposematism for gregarious larval survival, whilst identifying new questions about the roles of body size and toxicity in the evolution of grouping behavior.

## Introduction

Traits which, following classical Darwinism, must have evolved separately, and yet are highly beneficial when expressed together, raise questions about how, and in what order, such traits evolve. For example, social behavior often provides adaptive benefits through protection from predation, via various mechanisms from dilution to enhanced predator deterrence (Sillen-Tullberg 1988; Gamberale and Tullberg 1996a; Rowland et al. 2013; Svádová et al. 2014; Lehtonen and Jaatinen 2016), but group living often coincides with the evolution of other traits which serve similar purposes, such as physical and chemical defences (Hunter 2000; Nilsson and Forsman 2003; Beltran et al. 2007). Given this co-occurrence, one can ask whether associated traits, such as aggregation, promote the evolution of warning coloration, or whether warning signals facilitate gregarious behavior by providing protection that offsets the increased detection costs that grouping brings. This is a classic question about evolutionary sequence and causality involving two synergistic traits. However, in real populations associations are rarely dichotomous and a wider range of ecological and morphological traits may co-vary as adaptive suites of traits whose relationships are hard to unravel. Here, we focus on the impact of four key traits which are hypothesised to interact with the evolution of social behaviours: aposematism and structural defences, body size, and host plant use. We then test how some of these traits interact to aid larval survival in a natural setting.

Aposematism, the coupling of a conspicuous color pattern with a chemical defence (Ruxton et al. 2019), frequently co-occurs with gregariousness in larvae (Hunter 2000; Nilsson and Forsman 2003; Beltran et al. 2007). The evolution of toxicity in larvae is likely to be a strong driver for the evolution of warning signals and behaviors which amplify their effect, with toxicity widely believed to evolve before warning coloration and gregariousness (Sillen-Tullberg 1988; Alatalo and Mappes 1996; Tullberg and Hunter 1996; Ruxton and Sherratt 2006; Beltran et al. 2007; Boevé et al. 2018; Ruxton et al. 2019). Focus has therefore fallen on the causative links between aposematism and gregariousness, with plausible arguments presented for causality in both directions (see Ruxton and Sherratt 2006 for a review). First, protection from a chemical defence without a warning signal can be enough for gregariousness to be selected for (Alatalo and Mappes 1996; Riipi et al. 2001), yet by increasing detectability (and its associated costs), grouping may impose strong selection for the evolution of a conspicuous warning signal (Alatalo and Mappes 1996; Lindström et al. 1999; Riipi et al. 2001). Furthermore, a family grouping effect has long been proposed as a facilitator for the initial evolution of warning colours (Fisher 1930; Guilford 1985). However, evidence that aggregation is necessary for the evolution of aposematism is debated (Ruxton and Sherratt 2006; Dykema 2008). Aggregating without a warning signal to educate predators could be costly, suggesting instead that the prior acquisition of conspicuous warning coloration likely facilitates a switch to gregariousness (Sillen-Tullberg 1993; Tullberg and Hunter 1996; Tullberg et al. 2000; Wang et al. 2021). Recently, by re-analysing the seminal work of Tullberg and Hunter (1996), Wang *et al*. (2021) concluded that the latter scenario is more likely across Lepidoptera, but the uneven phylogenetic coverage of Tullberg and Hunter’s (1996) original dataset may influence some analyses, and the influence of other important traits is not explored. For example, morphological defences such as spines and thick hairs also deter predators (Greeney et al. 2012), but how these structural defences interact with gregariousness and aposematism remains unclear.

Larval body size is also likely to be critical to both the costs and benefits of gregariousness (García-Barros 2000; Nilsson and Forsman 2003). Aggregation may constrain body size via increased competition for food or by reducing the resources females can allocate to each egg in a cluster (Nilsson and Forsman 2003). Conversely, gregarious species may benefit from being larger if these constraints can be overcome, such as by enhancing the protection conferred by dilution with predators becoming satiated earlier when attacking a group (Curley et al. 2015). While the association between gregariousness and aposematism is well studied, as are the benefits that larger body sizes confer to signalling larvae (Gamberale and Tullberg 1996b; Forsman and Merilaita 1999; Nilsson and Forsman 2003; Mand et al. 2007; Sandre et al. 2007), much less is known of the three-way interactions between body size, aposematism and gregarious evolution. In a large phylogenetic comparative analysis of moth larvae, Nilsson and Forsman (2003) studied these three traits simultaneously but, due to statistical limitations at the time, were unable to consider all possible interactions between them, or pull apart their temporal patterns of acquisition. This leaves several important questions unanswered. For example, does larval body size influence whether gregariousness can evolve or not, perhaps through increased competition limiting resource acquisition or by delayed effects on adult female resource allocations? Or does gregarious behavior lead to selection for larger larval sizes, perhaps by amplifying an aposematic signal or offering the protection needed for longer development times? Finally, does a larger size facilitate the acquisition of a signal, given higher detectability of large prey, or vice versa?

A crucial factor in how the relative costs and benefits of larger body size play out is, in turn, the size and nature of the larval host plant. The size of the host plant, and therefore the amount of resources available to larvae, may dictate the level of competition for resources between individuals in a group (Tullberg and Hunter 1996; Nilsson and Forsman 2003). Indeed, previous phylogenetic analysis has revealed that some features of *Papilio* larval host plants, such as growth habit, leaf width and vegetation coverage, are important in the evolution of aposematism (Prudic et al. 2007; Gaitonde et al. 2018), but it is unclear whether the same is true for gregariousness. The spatial distribution of host plants, specifically whether they are patchy, may also influence whether adults lay their eggs in clusters and larvae develop gregariousness (Young 1983; Boevé et al. 2018).

Here, we use the diversity of butterfly larvae to address these questions as a case study in the interactions between behavioral and morphological traits. We systematically collected a new dataset of 353 species for the presence/absence of gregarious behavior and aposematic signals using a genus-level phylogeny of butterflies as a guide to ensure even phylogenetic coverage. We employ modern phylogenetic analyses to examine the evolution of larval gregariousness across butterflies, including the understudied roles that other related larval traits play in the acquisition of this behavior. As well as establishing coevolutionary interactions, we resolve uncertainty in key ecological transitions by determining the most likely causal pathways between traits, revealing which behaviors and morphologies make others more likely to evolve. We also test the assumptions of our phylogenetic models by investigating the effect that various combinations of traits have on the survival of artificial ‘caterpillars’ under avian predation in the field.

## Methods

### A. Phylogenetic analyses

#### i) Data collection

We obtained a list of 994 butterfly species, representing 991 genera and seven families, from a recently published phylogeny by Chazot *et al*. (2019). Using this list, we mined published literature for data on larval feeding behavioral status (solitary or gregarious) using Google Scholar. Where it was not specifically stated, we inferred solitariness from reports or images of adults laying eggs singly, and from visual evidence of leaf-rolling behavior, as was the case with many species in the Hesperiidae family. Although this approach may introduce some error when coding for solitariness, clustered egg laying and gregariousness are tightly correlated across species (Young 1983).

Once we had collected all available data on social behavior, we gathered information on four other traits of interest: i) larval color pattern (cryptic or aposematic), ii) the presence/absence of a structural defence (hairs, spines or tubercles), iii) final instar body length, and iv) the growth habit (or plant type) of the preferred larval host (Tables S1-S5; see Supplementary methods for details). To increase species coverage, adult wingspan (mm) was used as a proxy for final instar body length (c.f. Nilsson and Forsman 2003), which we validated by comparison with a smaller dataset on larval body length, finding a strong correlation between the median larval body length and log_10_ transformed adult wingspan (PGLS estimate = 0.007, SE = 0.001, *t* = 10.737, p < 0.001, n = 106). Four main online resources (Robinson et al. 2010; Warren et al. 2016; Kunte 2020; USDA 2021) were used to collect information on these traits. Larval warning coloration, in the form of bright, conspicuous colors coupled with contrasting dark colors was used to infer aposematism, as caterpillars with these signals are likely to be chemically defended (Tullberg and Hunter 1996). Finally, we recorded host plant data as categorical (graminoid, vine, forb/herb, shrub, tree) but converted these data into the binary trees (0) or not trees (1) for all analyses except in two regression models (indicated below). Trees were the most frequent type of host plant, with other categories individually too rare to be meaningfully analysed. In total, data on gregarious behavior were collected for 353 species from five families (Table S1), of which data were also available for color pattern in 290 species (Table S2), for structural defence in 237 species (Table S2), for final instar larval body length in 106 species, for adult wingspan in 261 species (Table S3), and for larval host plant type preference in 251 species (Table S4 & S5). 180 species had data for all five traits (excluding larval body length).

#### ii) Phylogenetic tree

The phylogenetic tree from Chazot *et al*. (2019, supplementary material S3b) was used for all phylogenetic analyses, which were performed in R v. 3.6.2 (R Core team 2013). Taxa with no data on social behavior were pruned along with their branches and internal nodes using the *ape* package (Paradis and Schliep 2018). The Lycaenidae were omitted from this study because many of the species within this family have highly derived ant associations, which is not captured in our focal traits and which has a strong influence on the evolution of their social behavior (Costa and Pierce 1997). All statistical analyses were performed in this phylogenetic framework.

#### iii) Quantitative analyses

We determined the phylogenetic signal of social behavior using Pagel’s λ (Pagel 1999). We then used the ‘fitDiscrete’ function in *geiger* (Pennell et al. 2014) to determine the most likely model by which transitions between character states have occurred. We used Akaike’s information criterion (AIC) to select between the null model, which assumes equal within-trait evolutionary transition rates between character states, with the all-rates-different model, which assumes the rate of transition from solitariness to gregariousness is different from the rate of transition from gregariousness to solitariness. Using the best-fitting model, we ran the ‘make.simmap’ function within *ape* (Paradis and Schliep 2018) to reconstruct the evolution of social behavior throughout the phylogeny, applying a maximum likelihood approach. This provides an estimation of the behavioural state (solitary/gregarious) at each node, which we used to determine the most likely ancestral character state and the number of instances of independent evolution of gregariousness. As the only continuous trait, we estimated the pattern of body size evolution by comparing (using a likelihood ratio test) the fit of three separate models: Brownian motion, Ornstein-Uhlenbeck and early burst (Blomberg et al. 2003).

Next, we used the ‘fitPagel’ function in *phytools* (Revell 2012) to test for coevolution between binary traits. Here, the dependent model assumes that the transition rate between states in trait A is influenced by the character state of trait B, whereas the null model assumes that the evolution of trait A is independent from trait B. We selected the best-fitting model based on AIC value, deferring to the simpler null model if there was less than two AIC points difference between models (Burnham and Anderson 2002). We then used MCMCglmm (Hadfield 2010) for detailed tests of co-evolution between traits and character states, whilst controlling for phylogenetic effects. We ran MCMC models for 5.1 million iterations, with a 0.1 million burn-in and sample storage frequency of every 500 iterations, with significance of the model calculated as the probability of the parameter value being different from zero (*P*_*MCMC*_).

Finally, we performed phylogenetic pathway analyses using the package *phylopath* (von Hardenberg and Gonzalez-Voyer 2013) to assess the order of trait acquisitions throughout the phylogeny. This method compares multiple potential evolutionary pathways to reveal which is most likely given the available tip data. Our first set of models included data on all traits (180 species) and considered the interactions between them simultaneously. Next, we focused on effects of body size in the gregarious-aposematism pathway using a larger dataset (229 species) and a simpler model set to consider these three traits specifically. Finally, we performed an additional analysis on an expanded dataset (217 species) to test our hypothesis that larval host plant preference directly influences the transition to gregariousness.

### B. Field experiments

#### i) Target details

To test how behavioral and morphological trait combinations may affect the survival of extant larvae in nature, we exposed artificial ‘caterpillar’ targets, that were either cryptic or aposematic, and grouped or solitary (figure 1A), to wild avian predation. In experiment 1, we tested these targets against green or white ‘leaf’ backgrounds so that we could distinguish differences in predation rate due to aversiveness versus detectability. Against white paper, both the cryptic and aposematic prey were highly detectable, whereas against green the cryptic prey was necessarily a better color match. In experiment 2, we varied the size of the targets (figure 1B) to assess the influence this had on predators’ responses. Our target sizes fell within normal the range of final instar body sizes seen across the butterfly phylogeny (figure 1C). We used a three-lobed shape for the ‘leaf’ platforms to roughly resemble the appearance of bramble leaves at their stem tip, with platforms subsequently being placed in locations with an abundance of bramble. Finally, to manipulate our targets’ palatability, dried mealworms (Wilko dried mealworms, Wilko Ltd, JK House, Notts, UK) were left to soak for no less than 24 hours in either plain water (for the cryptic/plain treatment) or a 2.5% Bitrex™ aqueous solution (5 g denatonium benzoate (Merck, Darmstadt, Germany) dissolved into 200 ml water, for the aposematic treatment). Plain mealworms were inserted into cryptic tubes and Bitrex™-soaked mealworms were inserted into aposematic tubes.

**Figure 1.**
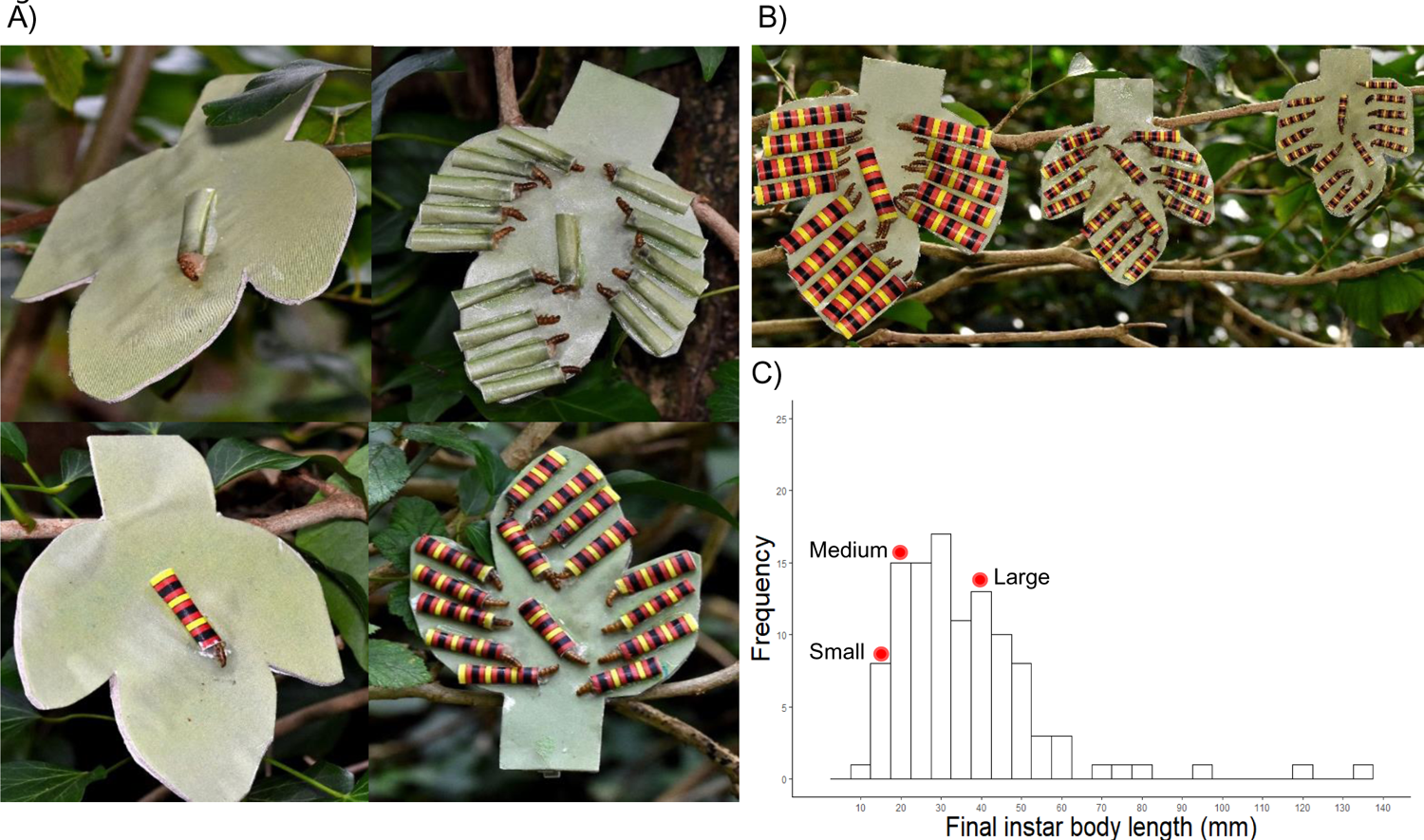
A) Examples of the artificial ‘caterpillars’ used in our field experiments on a green ‘leaf’ background. Clockwise from top-left: solitary cryptic, grouped cryptic, grouped aposematic and solitary aposematic. All targets are of ‘medium’ size. B) Examples of the three ‘body size’ treatments used in our second field experiment. From left to right: large, medium and small. C) The distribution of final instar larval body lengths from 106 species used in our analyses. Points along the x axis denote the sizes of the targets used in our field experiment.

We created eight treatments to be used in a 2 × 2 × 2 factorial design in the background color experiment. Treatments consisted of all eight possible combinations of either a cryptic or aposematic target which was solitary or grouped and on a green or white background. For the first experiment focusing on the interaction between color and group size, all targets were of ‘medium’ size. For the second experiment, integrating the effects of body size, we created 12 treatments to be used in a 2 × 2 × 3 factorial design. Treatments consisted of all 12 possible combinations of either a cryptic or aposematic target which was solitary or grouped and either small, medium or large in size. All backgrounds were green in this experiment. See Supplementary Material A for additional treatment creation details.

#### ii) Survival protocol

Both experiments were carried out from October – November 2021 around the Purdown estate (51.578462, −2.568755) in Bristol, UK. We selected experimental sites based on an abundance of varied foliage (e.g. bramble, bushes, trees etc.) on which ‘leaf’ platforms could be attached. To increase the likelihood that the predator population would be different between sites, the distance between separate sites was never less than c. 30 m. After assessing predator activity at each site using a sample of cryptic model larvae, we ‘pre-trained’ the community using cryptic and aposematic targets for 48 h (figure S1) before commencing experiment 1. Across both experiments, we placed the ‘leaf’ platforms in random order (determined by assigning each platform a number and assembling the sequence using a random number generator) in a rough circle around the site, with no two platforms closer than 5 m. Platforms were placed at various heights depending on the available vegetation (e.g. some attached to bramble close to the ground, some attached to tree branches overhead). We collected platforms after 24 h. After a minimum of two days, we repeated the experiment at the same site using a new random order of platforms. This meant that each treatment had two replicates per block. We ran 20 blocks per experiment for a total of 40 blocks across the study. We set up the body size experiment at the same sites as the background color experiment, often putting out the platforms for the former on the same day that we collected the platforms of the latter.

We recorded avian predation events as when the mealworm was mostly or completely missing from its tube. With ‘large’ targets, either one or both mealworms missing from a single target was considered a full predation event. Mealworms predated by mice were easily recognised by the presence of droppings on the platform and the tubes shredded into small scraps, and were treated as missing data.

#### iii) Data analyses

All analyses were performed in R v. 3.6.2 (R Core team 2013). Treatment interactions and their effect on target survival were analysed using the *glmer* function within the package *lme4*, with binomial error and a logit link function (Bates et al. 2015). We followed an established, stepwise model selection process (e.g. in McLellan et al. 2021) beginning with the initial saturated model containing individual platform ID and experimental block number as random effects, and target color (two levels: cryptic and aposematic), group size (two levels: solitary (1) and grouped (15)), background color (two levels: green and white) and body size (three levels: small, medium and large) as fixed effects. Block was not significant, so all subsequent models only contained platform ID as a random effect. Starting with the highest order interaction term between fixed effects, non-significant interactions were removed from the model and the next highest order interaction term was tested using the ‘drop1’ function to perform a Likelihood Ratio Test; we repeated this process to obtain the minimal adequate model containing only significant terms and, for interactions, their component main effects. Significant interaction terms were investigated by splitting the data based on one factor’s levels and continuing the model selection process on each subset of data.

## Results

### The evolution of gregarious larval behavior and associated traits across butterflies

Although phylogenetic signal of social behavior is high, and not significantly different from one (X^2^(1) = 1.831, λ = 0.947, p = 0.176), our model estimated a total of 49 transitions to gregariousness across the phylogeny (averaged estimate from 10,000 trees, figure 2 (but see figure S2 for the full social behaviour phylogeny)). Transition rates between behavioral states are unequal (X^2^(1) = 5.955, p = 0.015), revealing a faster rate of transition from gregariousness to solitariness than the reverse (solitary-gregarious = 0.006, gregarious-solitary = 0.020, Table S6), with the most probable ancestral character state estimated to be solitary (0.899 for solitary vs 0.101 for gregarious at the ancestral node).

**Figure 2.**
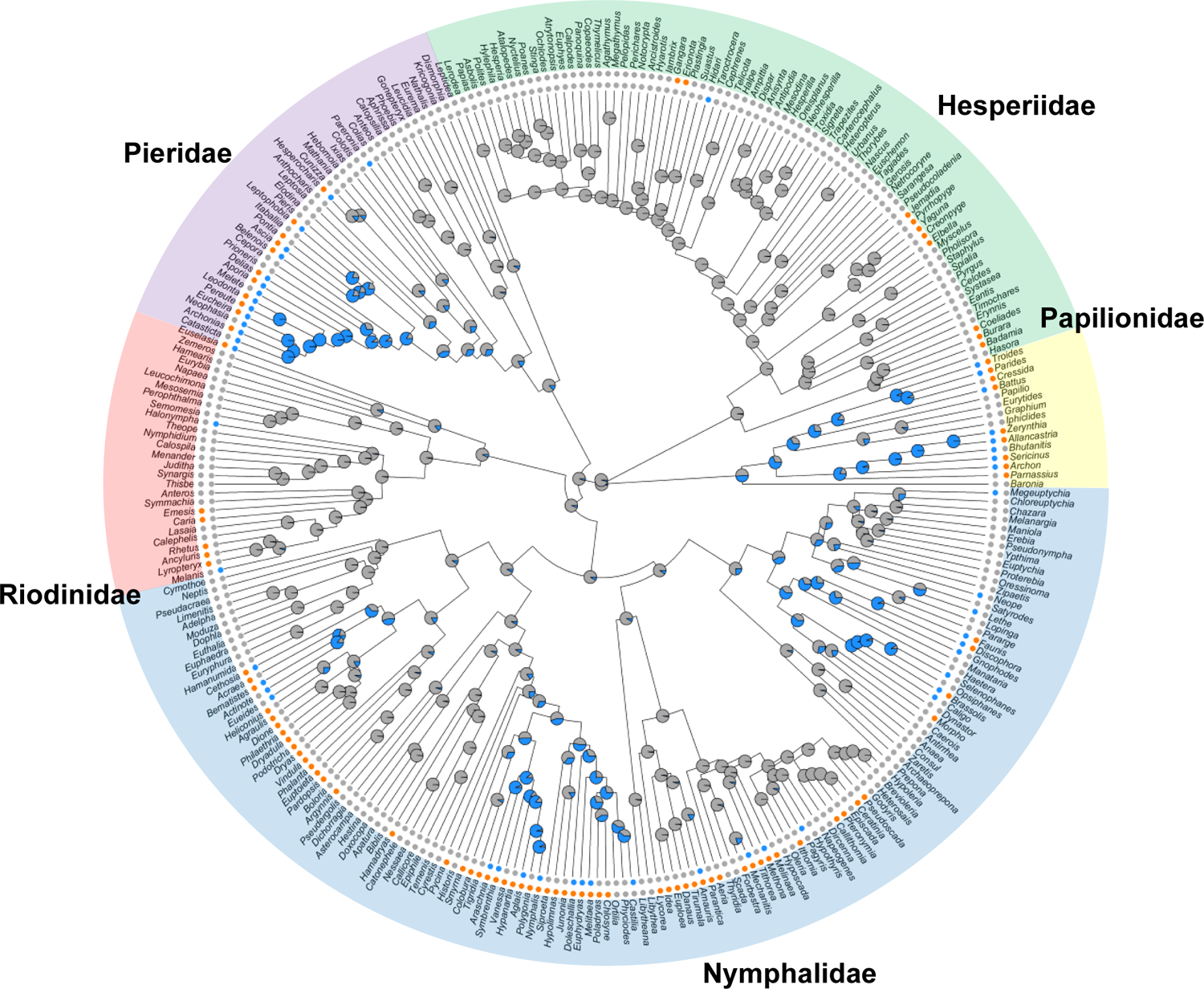
Genus-level butterfly phylogeny adapted from Chazot *et al*. (2019) showing estimated transitions to gregarious behavior (blue nodes and tips) from solitariness (grey nodes and tips). Aposematic (orange) and cryptic (grey) genera are shown in the outer ring of tips. The five separate families are highlighted

Traits putatively linked to larval gregarious behavior largely mirror the pattern of evolution of social behavior. The phylogenetic signal of color pattern is strong (X^2^(1) = 0.643, λ = 0.962, p = 0.423, figure 2), the rate of transition from aposematism to crypsis is faster than the reverse (model comparison: X^2^(1) = 11.241, p = 0.001; transition rates: cryptic-aposematic = 0.003, aposematic-cryptic = 0.016, Table S6) and crypsis is the estimated ancestral state (0.947 for crypsis vs 0.053 for aposematism). Similarly, the phylogenetic signal for structural defence evolution is high (X^2^(1) = 0, λ = 1, p = 1), the rate at which defences are gained and lost is equal throughout the phylogeny (model comparison: X^2^(1) = 0.015, p = 0.902; transition rate in either direction = 0.004, Table S6) and having no defence is ancestral (0.990 for no defence vs 0.010 for having a defence). Phylogenetic signal for body size is also high (X^2^ = 225.625, λ = 0.936, p < 0.001) and the most likely pattern of body size evolution follows a Brownian motion model (comparison with Ornstein-Uhlenbeck model: X^2^(1) = 3.127, p = 0.077; comparison with early burst model X^2^(1) = 0.001, p = 0.981), with the estimated ancestral adult wingspan to be 52.8 mm. Finally, larval host plant preference shows a strong phylogenetic signal (X^2^(1) = 2.844, λ = 0.914, p = 0.092) and equal transition rates between feeding on trees and other plant types (model comparison: X^2^(1) = 0.319, p = 0.572; transition rate in either direction = 0.011, Table S6). The equal rates model did not estimate the ancestral trait in favour of one state over the other (0.497 for trees vs 0.503 for non-trees but see Supplementary results).

### Coevolution between gregariousness and aposematism

We found that transitions from solitariness to gregariousness occur at a higher rate in aposematic lineages, whereas the reverse occurs at a higher rate in cryptic lineages (ΔAIC = 18.936, n = 290, figure 3A). Similarly, evolutionary switches from crypsis to aposematism are faster in gregarious lineages, whereas the reverse switch occurs more readily in solitary lineages (ΔAIC = 18.327, n = 290, figure 3A), revealing a clear coevolutionary relationship between the two traits. Solitary to gregarious transitions occur at a higher rate in lineages possessing a structural defence, whereas transitions away from gregariousness are faster in undefended lineages (ΔAIC = 2.884, n = 237, Table S7). The transition rate from crypsis to aposematism is faster in defended lineages and the reverse is faster without a defence (ΔAIC = 12.002, n = 237, Table S7). The rate of acquisition of a structural defence is higher in gregarious lineages and the rate at which defences are lost is higher in solitary lineages (ΔAIC = 7.671, n = 237, Table S7). The acquisition of a structural defence occurs faster in aposematic lineages, whereas losing a defence happens faster when cryptic (ΔAIC = 19.619, n = 237, Table S7).

**Figure 3.**
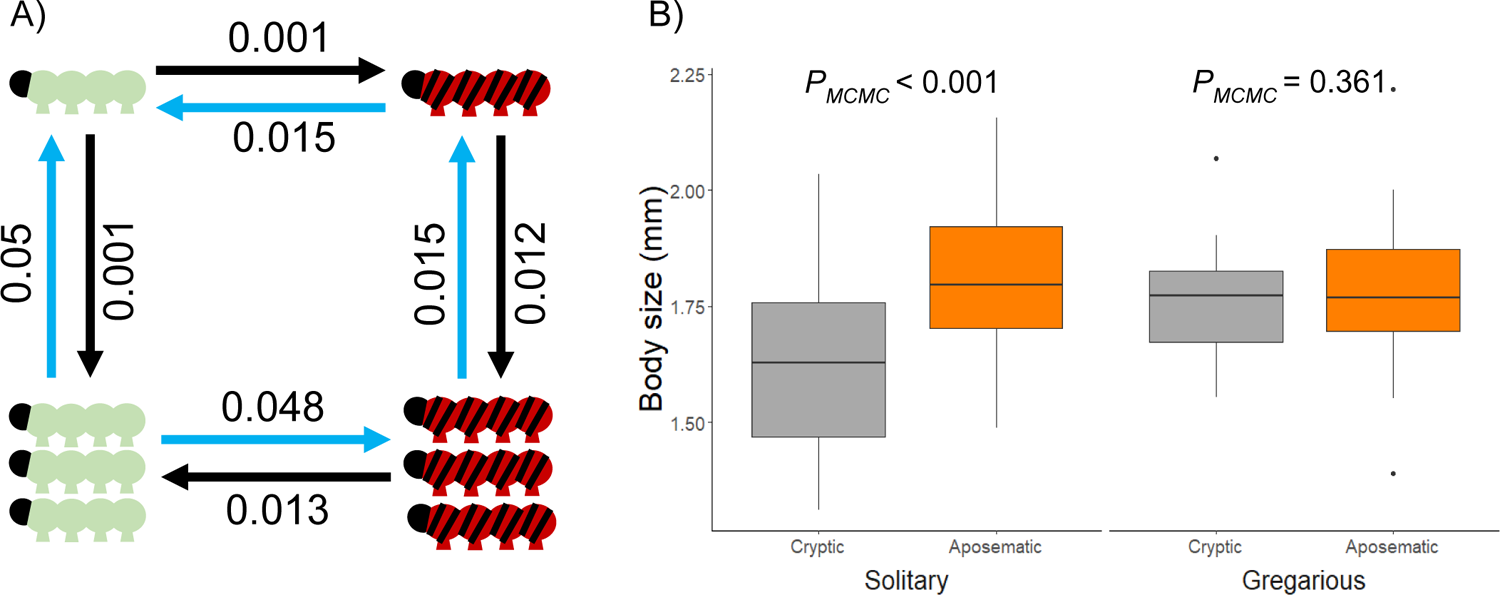
A) The results from two separate models looking at transition rates between character states of larval traits: solitariness and gregariousness depicted as one or three caterpillars respectively; crypsis and aposematism depicted as green or red/black striped caterpillars respectively. Blue arrows denote the higher transition rate of either direction. B) Comparisons of final instar larval body size between caterpillar species using different color pattern strategies. Aposematic (orange boxes) species are generally larger than cryptic (grey boxes) species most obviously when both are solitary.

Given the clear evidence of an evolutionary relationship between gregariousness and aposematism, we performed MCMCglmm regressions to examine this interaction in more detail. We first included all traits and interactions in the model, the final model after the simplification process revealed no significant interactions between factors so we ran simpler, two-factor models with larger datasets. A simple model with social behavior and color pattern revealed that gregarious species are more likely to be aposematic than cryptic (p-mean = 35.683, 95% CI = 2.166-71.316, *P*_*MCMC*_ = 0.009). We next examined structural defences as another morphological larval defence that might influence the evolution of gregariousness. Separate pairwise models revealed no interaction between social behavior and structural defences (p-mean = 56.785, 95% CI = −4.716-127.235, *P*_*MCMC*_ = 0.053), whereas aposematic species are much more likely to possess a defence than not (p-mean = 95.983, 95% CI = 5.589-236.509, *P*_*MCMC*_ < 0.001). The interaction between color pattern and structural defence does not influence the evolution of social behavior (p-mean = −10.98, 95% CI = −152.95-118.39, *P*_*MCMC*_ = 0.867). Additionally, two-factor tests for interactions between social behavior and categorical host plant, and color pattern and categorical host plant revealed no interactions between these traits and separate host plant types (see Supplementary results).

### Associations between larval body size, coloration and social behavior

There is no compelling evidence that the evolution of gregariousness is influenced by the interaction between color pattern and body size (p-mean = −89.43, 95% CI = −413.55-163.56, *P*_*MCMC*_ = 0.445) nor, after removing the non-significant interaction, is there a main effect of body size on gregariousness (p-mean = 80.77, 95% CI = −55.35-273.09, *P*_*MCMC*_ = 0.221). However, aposematic species are generally larger than cryptic species (p-mean = 131.83, 95% CI = 15.05-251.76, *P*_*MCMC*_ < 0.001). Despite the lack of statistical evidence for an interaction, the body size difference seems more apparent in solitary species (solitary species only: p-mean = 211.26, 95% CI = 22.91-474.41, *P*_*MCMC*_ < 0.001, which remains significant after Bonferroni correction for two unplanned comparisons; figure 3B) but not gregarious species (gregarious species only: p-mean = 99.75, 95% CI = −147.20-430.47, *P*_*MCMC*_ = 0.361; figure 3B).

### Trait evolutionary pathways

After examining how different traits are associated with one another, we explored the causal relationships between traits by testing whether the acquisition of a given trait makes the subsequent evolution of others more likely. Given the close relationship between gregariousness and aposematism revealed by our phylogenetic regression analyses, we predicted a direct causal relationship between these two traits. Our initial full model (n = 176) showed strong support for the pathway in which, starting with a solitary, cryptic and undefended ancestor, a structural defence is first acquired, which leads to the evolution of a warning signal (pathway coefficient = 0.900), which in turn facilitates the evolution of gregarious behavior (pathway coefficient = 0.923, figure 4A).

**Figure 4.**
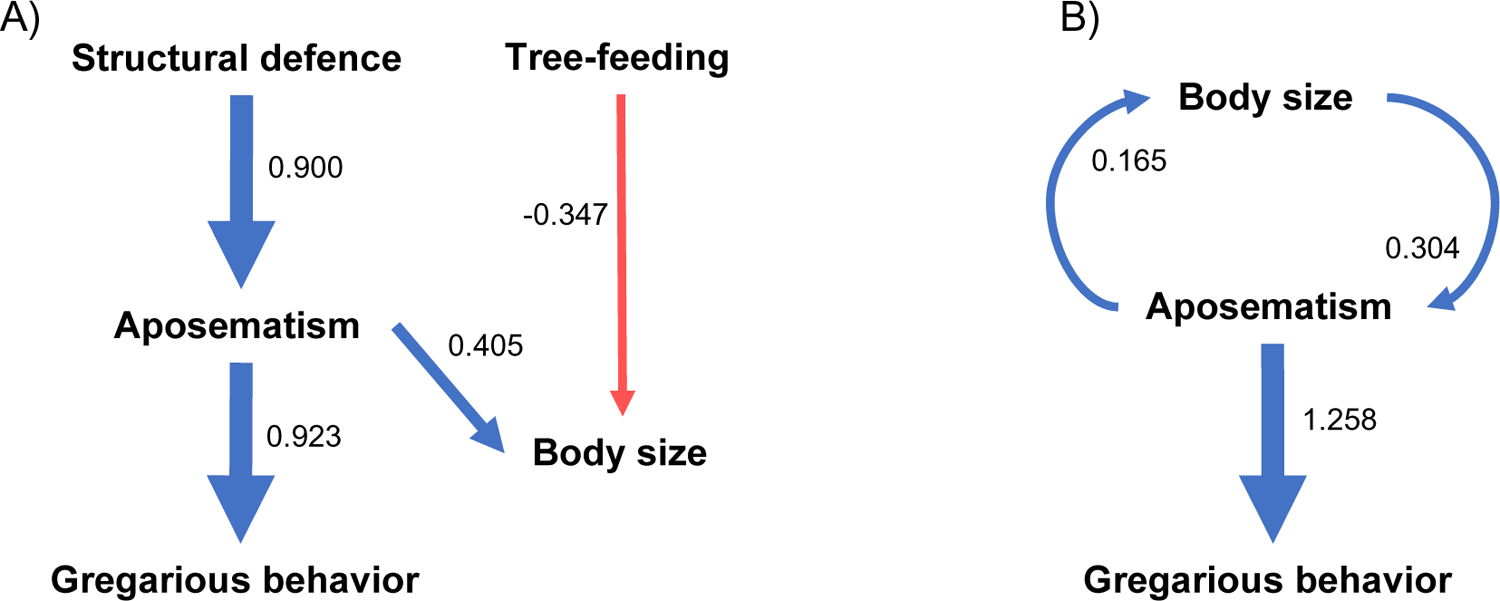
Results from phylogenetic pathway analyses (PPA) in which evolutionary causal relationships between traits are estimated. Arrows show the direction of causal evolutions, the values next to arrows display the pathway coefficient (strength of the estimated relationship). Positive coefficients (blue arrows) indicate that a 0-1 transition in the parent trait predicts a 0-1 transition in the child trait (or an increase in the case of the continuous body size trait). Negative coefficients (red arrow) indicate that a 0-1 or 1-0 transition in the parent trait predicts a respective decrease or increase in the child trait value. A) Output from the saturated model showing strong support for the pathway in which an initial acquisition of a structural defence predicts a transition to aposematism, which in turn predicts the adoption of gregarious behavior. B) Output from the simple three-trait model showing equal support for an increase in larval body size predicting a switch to aposematism or vice versa.

Our second, larger model (n = 229) focused on the interactions between social behavior, color pattern and body size, and suggests a causal relationship between larval body size and color pattern in both directions (figure 4B). One possible interpretation of this model is that an initial evolution towards a larger body is causative in promoting the acquisition of a warning signal (pathway coefficient = 0.304, model CICc = 13.0), and this signal then promotes gregariousness (pathway coefficient = 1.258). Alternatively, the evolution of aposematism may promote an increase in body size (pathway coefficient = 0.165, model CICc = 13.0), and/or the subsequent evolution of gregariousness (pathway coefficient = 1.258). The model does not support a direct relationship, in either direction, between larval body size and social behavior.

### Experimental evidence for interactions between group size, colors and body size

Data from experiment 1 revealed that the overall three-way interaction between target color, group size and background color was significant (X^2^(1) = 7.718, *p* = 0.005), therefore we explored the two-way interactions. The interaction between background color and group size did not affect survival of cryptic targets (X^2^(1) = 0.026, *p* = 0.872), but did influence aposematic target survival (X^2^(1) = 8.722, *p* = 0.003). Background color had no effect on solitary aposematic target survival (X^2^(1) = 0.054, *p* = 0.816), whereas grouped aposematic targets survived better when on a green background (X^2^(1) = 5.868, *p* = 0.015; Table S8) and had the highest survival (∼15% higher than the second-best surviving treatment) across experiment 1 (Table S8).

In the second experiment, incorporating body size variation, the overall three-way interaction between target color, group size and body size was not significant (X^2^(2) = 0.322, *p* = 0.851), we therefore removed non-significant terms in a stepwise manner to reveal any remaining significant terms. The interaction between target color and group size influenced target survival (X^2^(1) = 61.714, *p* < 0.001), where cryptic, solitary targets survived better than cryptic, grouped targets (X^2^(1) = 98.234, *p* < 0.001), and aposematic grouped targets survived better than aposematic solitary targets (X^2^(1) = 4.738, *p* = 0.030). Target survival was not affected by the interactions between target color and body size (X^2^(2) = 0.325, *p* = 0.850) and group size and body size (X^2^(2) = 0.742, *p* = 0.690). However, we note the non-significant trend towards both cryptic and aposematic targets surviving comparatively worse when large, with a marginally steeper change in solitary targets (figure 5).

**Figure 5.**
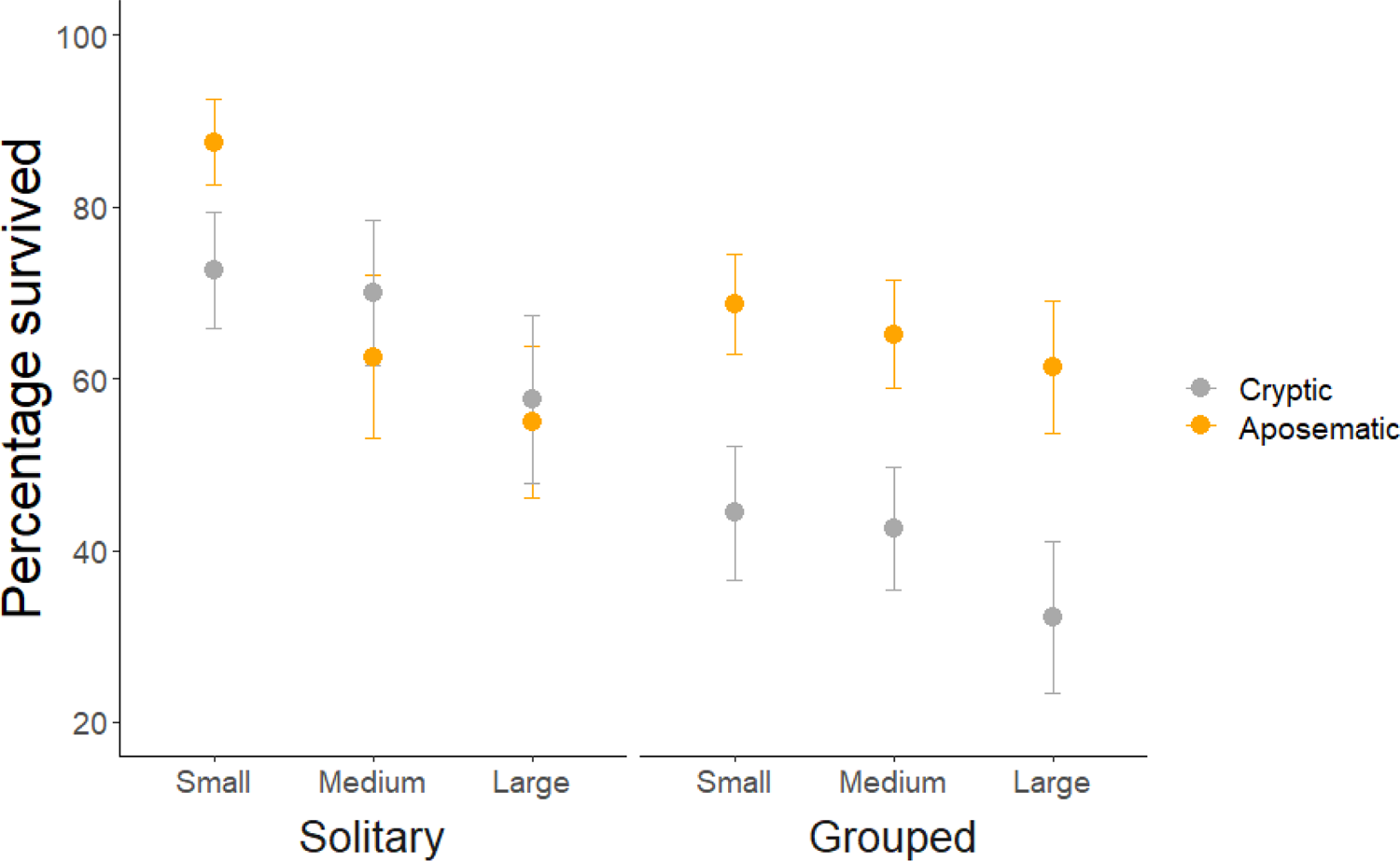
A comparison of treatment survival between different ‘body sizes’ across our second field experiment. Cryptic target data are displayed with grey points and aposematic target data with orange points. Error bars show 95% CI. Size did not have a significant effect on survival overall, but we note the negative trend in survival with increasing body size in both group size and color pattern treatments. Aposematic targets survived better than cryptic targets at all sizes when both were grouped.

To examine the overall interaction between target color and group size, we combined the green background and medium body size data from both experiments to gain statistical power. We found an overall interaction between target color and group size (X^2^(1) = 21.286, *p* < 0.001), where cryptic targets survived better when solitary (X^2^(1) = 6.704, *p* = 0.010), whereas aposematic targets survived better when grouped (X^2^(1) = 6.644, *p* = 0.010; figure 6). Additionally, there was no difference in survival between solitary targets of either color (X^2^(1) = 0.051, *p* = 0.821).

**Figure 6.**
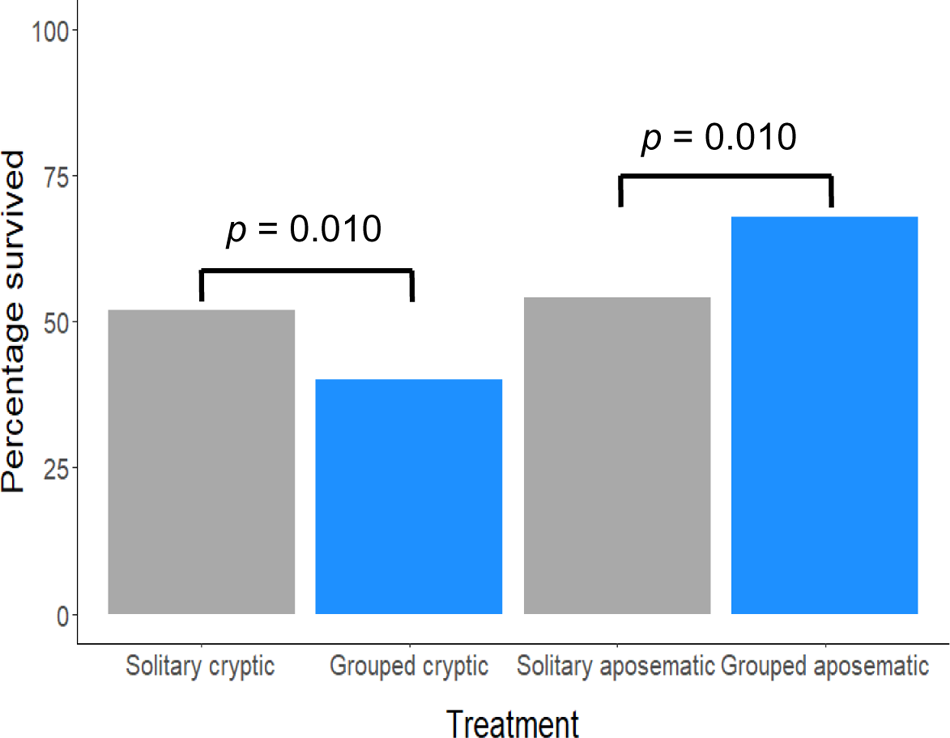
Comparing treatment survival using pooled data from both field experiments, only ‘medium’ sized, green background targets were considered. Grouped targets (blue bars) survived worse than solitary (grey bars) when cryptic, but better than solitary when aposematic.

## Discussion

Our results show that, from a solitary, cryptic ancestor, gregariousness has evolved many times across the phylogeny, but with relatively low rates of occurrence among extant species. This contrast between repeated evolution and low occurrence suggests that gregarious behavior is highly evolvable, but must have considerable costs, with the conditions which facilitate its evolution and maintenance being met relatively infrequently. It is likely that these favourable conditions include the presence of other, complementary traits. We show here that aposematism is highly likely to be a key facilitator in the evolution of gregariousness, more so than vice versa. Our comparative data also reveal an evolutionary link between final instar body size and warning coloration, where solitary aposematic species are, on average, larger than cryptic species.

Why is aposematism a predisposition for gregariousness? While grouping behavior provides protection via the dilution effect (Sillen-Tullberg 1988; Lehtonen and Jaatinen 2016), an effective defence can be bolstered by aposematism (Hunter 2000; Beatty et al. 2005; Curley et al. 2015). In turn, aggregation improves the anti-predator effect of warning signals (Gamberale and Tullberg 1996a; Rowland et al. 2013; Svádová et al. 2014) and can increase the speed of predator avoidance learning (Harvey et al. 1982; Riipi et al. 2001; Rowland et al. 2013; Svádová et al. 2014). Other recent work also suggests that a warning signal is most likely to evolve before, and subsequently facilitate the evolution of, gregarious behavior (Wang et al. 2021). Our analyses of a novel phylogenetic dataset support the conclusion that, in most cases, aposematism favours the evolution of gregarious behavior, whereas aggregation is not a necessary prerequisite for the evolution of a warning signal (Sillen-Tullberg 1993; Tullberg and Hunter 1996; Tullberg et al. 2000; Ruxton and Sherratt 2006; Dykema 2008). The evolutionary pathway supported here likely reflects the disadvantages that larvae face by aggregating without a warning signal to dissuade predators (Mappes and Alatalo 1997), as evidenced in our field data. Our experimental data show that cryptic and undefended ‘caterpillars’ survive better when solitary, regardless of their body size, most likely because of the high detectability costs that come with aggregating (Riipi et al. 2001). This suggests that, even with near-perfect color match to the background, aggregating when undefended puts larvae at a relative disadvantage. Conversely, our aposematic targets survived better when grouped, also at each body size, suggesting aposematism provides the protection and reduction in detectability costs that allows gregariousness to evolve. For solitary larvae, our field data indicate that crypsis and aposematism are equally viable strategies, which supports the evolutionary pathway in which warning coloration can arise in solitary species relatively easily. We were also interested in how conspicuity against the background affects larval survival. That aposematic grouped targets survived best against a green background is difficult to interpret, given we assume the white background made all targets more conspicuous which should particularly effect cryptic prey. It is possible that more visible white background attracted more naïve predators, or the targets and/or the platform itself were slightly less detectable with a green background, such that the combination of lower detection and predator acceptance benefits aposematic groups on green backgrounds, but this does not explain the lack of difference in predation between other treatments.

We also found that aposematic larvae are likely to possess a structural defence, and that the acquisition of such defences likely precedes the evolution of aposematism, supporting previous theoretical work which indicates that spines can evolve in larvae before warning coloration (Speed and Ruxton 2005). This result can be explained in terms of detectability. Solitary, cryptic larvae evolve dense hairs or spines as protection against invertebrate and/or vertebrate predation (Greeney et al. 2012). These defences might increase detectability (Speed and Ruxton 2005), making crypsis a less effective strategy, thus favouring the evolution of a conspicuous signal. The increased detectability costs conferred by grouping is then easier for signalling species to acquire, as gregariousness may also enhance the protection offered by a structural defence (Hunter 2000).

Our results also provide new evidence of interactions between body size and traits impacting detectability. In contrast to previous suggestions that competition for food between group members will constrain larval growth (Nilsson and Forsman 2003), we did not observe an overall interaction between final instar larval body size and social behavior. However, within the subset of solitary species, we found that aposematic larvae are generally larger-bodied than cryptic species (figure 3B). This finding supports previous observations that the evolution of aposematism is associated with larger body sizes (e.g. in Medina et al. 2020). This link is likely explained by the enhanced anti-predator effect afforded to larger aposematic signals (Gamberale and Tullberg 1996b; Forsman and Merilaita 1999; Nilsson and Forsman 2003; Mänd et al. 2007). We did not find evidence of a similar evolutionary interaction between color pattern and body size across gregarious larvae, suggesting there is only strong selection for solitary species to present larger signals. This is probably because solitary individuals can only display a larger signal by increasing their body size, whereas aposematic larvae that aggregate can collectively augment their displayed signal to be more aversive to predators (Gamberale and Tullberg 1996a; Mappes and Alatalo 1997; Gamberale-Stille 2000; Wang et al. 2021). However, we did not observe a positive interaction between body size and survival of aposematic targets in our field experiments, although we observe a slight trend towards large aposematic targets surviving worse than their smaller counterparts, especially when solitary. This trend is more consistent with evidence that remaining undetected is a better strategy for aposematic larvae than a maximally conspicuous signal, as suggested in various studies of distance-dependent camouflage (Barnett and Cuthill 2014; Barnett et al. 2016), and may warrant further investigation. For example, it may simply be the case that our chosen selection of ‘body sizes’, representing incremental increases between groups, was not sufficient to elicit differences in response from predators, and greater effects could be found with larger prey.

The association between aposematism and body size is established in the literature, yet the causative evolutionary relationship between these traits has previously gone unstudied. Our data present two equally likely scenarios which principally differ in the order in which aposematism and an evolutionary increase in body size occur. In the first scenario, selection favours a larger body in solitary, cryptic larvae, which subsequently promotes the acquisition of a warning signal, making a switch to gregariousness more likely. Benefits to adult fecundity might result in a drive towards larger sizes in cryptic insects (Honěk 1993; Mänd et al. 2007; Remmel and Tammaru 2009), and the higher detectability costs experienced by larger prey should allow warning coloration to evolve more readily than in smaller species (Nilsson and Forsman 2003). Results from our field experiments give some (non-significant) indication that there may be predator-related costs to larger body sizes for cryptic, undefended larvae (figure 5). These costs, if they exist, would most likely be the result of increased detectability (Riipi et al. 2001; Remmel and Tammaru 2009), as evidenced by the improved survival of cryptic solitary targets compared to grouped, regardless of body size. Thus, both our comparative and field data suggest that beyond a certain level, detectability costs of increased body size render crypsis an ineffective strategy for protecting larvae. Instead, conspicuous signalling (assuming a prior toxic defence) is likely the better strategy at larger sizes. Crypsis may be favoured before these larger sizes if warning signals are less effective as deterrents while maintaining their production costs (Mänd et al. 2007), although we did not observe this effect in our current field data. Whether this shift in effective strategy exists, and at what relative size it might occur, merits further investigation.

In the second scenario, the initial acquisition of a warning signal promotes a subsequent increase in body size, and simultaneously makes gregarious behavior more likely to evolve. Signals displayed by small larvae may not be very effective at deterring predators (Mänd et al. 2007; Hossie et al. 2015), therefore the evolution of a warning signal should apply considerable pressure on prey to increase in size (Forsman and Merilaita 1999; Nilsson and Forsman 2003), and could facilitate longer larval development times (Speed and Ruxton 2005). Our field data did not reveal a survival disadvantage for our small, aposematic targets. However, it is possible that the target’s signal was large enough to be aversive to birds across all body sizes. Additionally, our small targets were just as unpalatable as the larger targets, so birds may have rejected them at the same rate. Conversely, larvae in nature often have a lower toxin load during their early instars, compared to when they are older and larger (Merilaita and Tullberg 2005; Grant 2007), and thus are at greater risk of predation.

In summary, the evidence that aposematism arises before, and facilitates, gregarious behavior, and that aggregating is in fact detrimental for undefended larvae is clear from our study. Gregarious larvae gain considerable anti-predator benefits from aposematism, which in most cases appears important for aggregated larval survival. Conversely, solitary larvae benefit equally from a cryptic or aposematic strategy. Although it is clear from our data that aposematism promotes a switch to gregarious behavior, other factors which make this transition favourable over remaining solitary require further investigation. Larval body size and aposematism appear closely linked evolutionarily from our phylogenetic data, although we found no evidence that these traits directly interact to improve larval survival in a natural field setting. It is possible that solitary larvae are under selection for a larger body both when cryptic and aposematic. In the former case, there may be a body size above which a switch to aposematism becomes increasingly favourable. We note that the acquisition of a chemical defence during larval butterfly evolution is an important component missing from our current study. In our analyses we assume aposematism is an accurate proxy for toxicity, but the potential for independent evolution of toxicity likely has a considerable impact on the acquisition of the traits we have considered. For example, does a switch to a toxic host plant (and thus the acquisition of a toxic defence) in solitary, cryptic larvae trigger an increase in body size, as has been theorised for grasshoppers (Whitman and Vincent 2008)? Is there strong selection to acquire a chemical defence in large, cryptic larvae? Or does selection act on all solitary and cryptic species to grow larger, but only the ones that switch to toxic hosts develop warning signals? Toxicity is widely believed to evolve before warning coloration and gregariousness (Sillen-Tullberg 1988; Alatalo and Mappes 1996; Tullberg and Hunter 1996; Ruxton and Sherratt 2006; Beltran et al. 2007; Boevé et al. 2018; Ruxton et al. 2019), as these traits would be largely detrimental to larval survival without it. This evolutionary pathway is mirrored by the ontogenetic changes of the desert locust (*Schistocerca gregaria*), where a behavioral switch to feeding on toxic hosts precedes both gregariousness and aposematism (Despland and Simpson 2005). The evolution of toxicity in relation to body size and structural defences presents an interesting topic for future phylogenetic investigation. However, our present findings go some way to explain the interactions between larval body size, coloration and social behavior once toxicity has evolved.

## Supporting information

Supplemental R scripts

## Ethics statement

Bitrex™ is non-toxic and does not harm birds when ingested. We received approval from the Animal Welfare and Ethical Review Body (AWERB) of the University of Bristol to conduct our field study.

## Acknowledgments

Many thanks to Dr Kevin Arbuckle for his help and advice with some phylogenetic analysis and interpretation. Thanks also to members of the EBaB lab group for help with target creation.

## Author contribution statement

S.H.M. conceived the research; S.H.M., I.C.C. and C.F.M. designed the experiments; C.F.M. carried out the data collection and field experiments; S.H.M. and I.C.C. assisted C.F.M. with data analyses and C.F.M. wrote the first draft of the manuscript with subsequent contributions by all authors.

