## Supplemental material for "Warning coloration, body size and the evolution of gregarious behavior in butterfly larvae"

**Supplementary tables (see separate excel spreadsheet ‘Larval_traits_references’):**

- **Table S1:** Recorded data for larval social behavior with literature citations. The ‘Social behaviour from lit’ column displays all instances of reported social behaviour for a given species, where ‘Sol’ = solitary, ‘Greg’ = gregarious and ‘Chng’ = social behaviour changes throughout larval development. The ‘Social behaviour in data’ column represents how the data were recorded for analyses, where 0 = solitary and 1 = gregarious. Any recorded instance of gregarious behaviour within a genus (represented by ‘spp’ in the ‘Species’ column) or during larval development (represented by ‘Chng’ in the ‘Social behaviour from lit’ column) are recorded as 1 in the data.
- **Table S2:** Recorded data for larval social behavior, color pattern and structural defence with literature citations. Social behavior recorded as in Table S1 except ‘Chng’ entries from literature recorded in data as solitary. Color pattern recorded as 0 = cryptic and 1 = aposematic. Structural defence recorded as 0 = no defence and 1 = has defence.
- **Table S3:** Recorded data for larval social behavior, final instar larval body length and adult wingspan with literature citations. Social behavior recorded as in Table S2 ‘Social behaviour in data’ column. Both larval body size and wingspan columns show the range of sizes recorded from the literature
- **Table S4:** Recorded data for larval social behavior, larval host plant genus, species and plant species growth habit with literature citations. Social behavior recorded as in Table S2 ‘Social behaviour in data’ column.
- **Table S5:** All recorded growth habits for larval host plant genera as recorded in Table S4 with literature citations.

**Supplementary methods**

**Host plant data collection**

Host plant data were collected by first performing a species-level search of larvae within the Natural History Museum’s (NHM) HOSTS webpage (Robinson *et al*., 2010) and recording every host plant genus, and some species, that a given species was reported to feed on. Alternative literature sources were used for species not included in the NHM HOSTS database. We then recorded the described growth habits of the plant genera mainly within the USDA PLANTS database (USDA, 2021), wherein entries listed as ‘subshrub’ were included as ‘shrub’. Plant species with multiple recorded growth habits were refined to a single growth habit on the basis of whatever the most frequent (mode) recorded growth habit was. Where the mode was equal between two or more growth habits, a single entry was decided based on whichever was assumed to be the larger growth habit. Size rankings were: graminoid < vine < forb/herb < shrub < tree.

**Target creation**

The green color used for the ‘leaf’ background and cryptic target treatment was created from calibrated photographs of 50 bramble leaves, mapped to the colour space of a typical insectivorous passerine, the blue tit (Cyanistes caeruleus) (Hart *et al*., 2000). Different green shades were printed iteratively until a close match to mean 'bramble green' was found (see Barnett *et al*., 2018 for further details). The warning signal colors and pattern were selected based on typical aposematic displays (Stevens and Ruxton, 2012), without mimicking any species that local birds are likely to have encountered in the wild. This was to avoid predators’ prior experience confounding target predation rates. We printed color patterns onto waterproof paper (Rite-in-the-Rain, J.L. Darling LLC, Tacoma, WA, USA) and rolled them into hollow tubes, kept together with glue (Wilko all purpose adhesive, Wilko Ltd, JK House, Notts, UK). Targets were rolled into ‘small’ (15.92 mm length, 2 mm diameter), ‘medium’ (22 mm length, 3.4 mm diameter) or ‘large’ (41.58 mm length, 7 mm diameter) cylinders, which roughly relate to the lengths of second, third and fifth instar *Pieris brassicae* larvae respectively (Bhowmik and Gupta, 2017). Using the medium target as a template, the length of the aposematic target was adjusted so that the separate elements of its color pattern remained in proportion relative to its size.

When necessary, mealworms were held in place with a small lump of dough (a mixture of plain flour, margarine (Stork vegan baking block, Stork, London, UK) and water). We left the head and thorax of the mealworm protruding from the tube, this allowed for easy identification of a predation event. For the body size experiment, mealworms were pre-sorted by eye into ‘large’ or ‘small’ categories. Large mealworms were approximately twice the length and/or diameter of small mealworms. Only small mealworms were inserted into small targets, large mealworms were inserted into medium targets and two large mealworms were inserted into each large target.

To create the ‘leaf’ platforms, we glued a sheet of waterproof paper to either side of a sheet of card (1500 micron craft board, House of Card and Paper, Amazon.co.uk). For green platforms, both sheets were green. For white platforms, one sheet was white while the underside was green, this was to ensure that only the background of the targets was different between treatments. A small clothes peg (clear mini pegs, WDAFLG, Amazon.co.uk) was glued to the underside of each platform, allowing it to be attached to foliage. In our body size experiment, the size of the ‘leaf’ platforms, but not the shape, was altered to accommodate each target size, so that the space between targets and their positioning was roughly equivalent to that of the medium targets.

**Survival protocol**

At each site, to gauge the level of predator activity, and give local predators an opportunity to learn the difference in palatability between our cryptic and aposematic targets, we set out ‘pre-training’ plates, consisting of individual targets glued directly onto palm leaf plates (20x20 cm disposable palm leaf plate, BIOZOYG, Amazon.co.uk) approximately 2 cm apart from one another. For the ‘cryptic’ pre-training plate, 15 cryptic targets were presented. For the ‘mixed’ plate, seven cryptic and eight aposematic targets were randomly distributed across the plate. Plates sat atop a single bamboo pole.

**Supplementary results**

When host plant was analysed as a categorical variable, the most likely ancestral state was estimated to be forb/herb-feeding (0.645 for forb/herb vs next closest 0.342 for tree). We also tested for a coevolutionary relationship between larval social behavior and (binary) host plant preference. Transitions between social behaviors occur independently of host plant preference (ΔAIC = -1.938, n = 251), and vice versa (ΔAIC = -2.872, n = 251), throughout the phylogeny. As the fitpagel function does not allow categorical variables to be included in the model, we performed an MCMCglmm regression on social behavior and host plant as a categorical variable to test for coevolutionary relationships on a finer scale, yet found no significant interaction terms (forb/herb: p-mean = -58.941, 95% CI = -203.681-51.962, *P_MCMC_* = 0.282; graminoid: p-mean = -46.836, 95% CI = -157.977-42.932, *P_MCMC_* = 0.281; shrub: p-mean = -40.128, 95% CI = -107.467-20.853, *P_MCMC_* = 0.162; tree: p-mean = 5.085, 95% CI = -49.688-64.250, *P_MCMC_* = 0.848; vine: p-mean = -52.116, 95% CI = -154.300-20.073, *P_MCMC_* = 0.111). A two-factor regression model between color pattern and categorical host plant also revealed no significant interaction terms (forb/herb: p-mean = -47.34, 95% CI = -175.17-62.41, *P_MCMC_* = 0.366; graminoid: p-mean = -82.11, 95% CI = -261.81-23.36, *P_MCMC_* = 0.133; shrub: p-mean = 11.84, 95% CI = -49.72-76.51, *P_MCMC_* = 0.696; tree: p-mean = -16.45, 95% CI = -74.39-36.57, *P_MCMC_* = 0.555; vine: p-mean = 27.07, 95% CI = -48.96-113.82, *P_MCMC_* = 0.479).

**Supplementary figures**

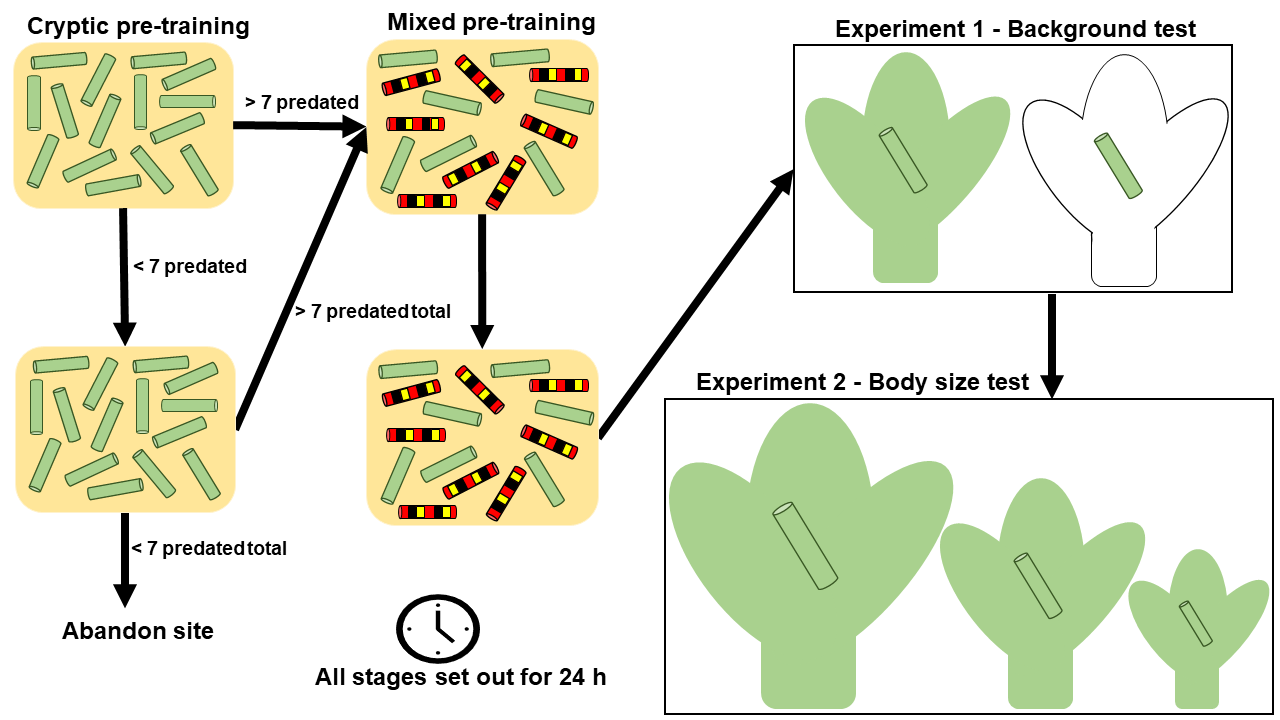

**Figure S1**

The order in which we conducted our field experiments. Starting from top-left, 15 cryptic targets were set out for 24 h. If fewer than seven targets were predated, we replaced the ‘cryptic’ pre-training plate with a fresh one to be left for another 24 h. If after 48 h fewer than seven targets were predated in total, we abandoned the site for the rest of the study. If, after 24 or 48 h, seven or more targets were predated in total, we replaced the ‘cryptic’ plate with a ‘mixed’ plate, which we replaced with another, fresh mixed plate after 24 h regardless of predation activity. We left the second mixed plate out for 24 h, after which we collected it and immediately began experiment 1, followed by experiment 2 the next day.

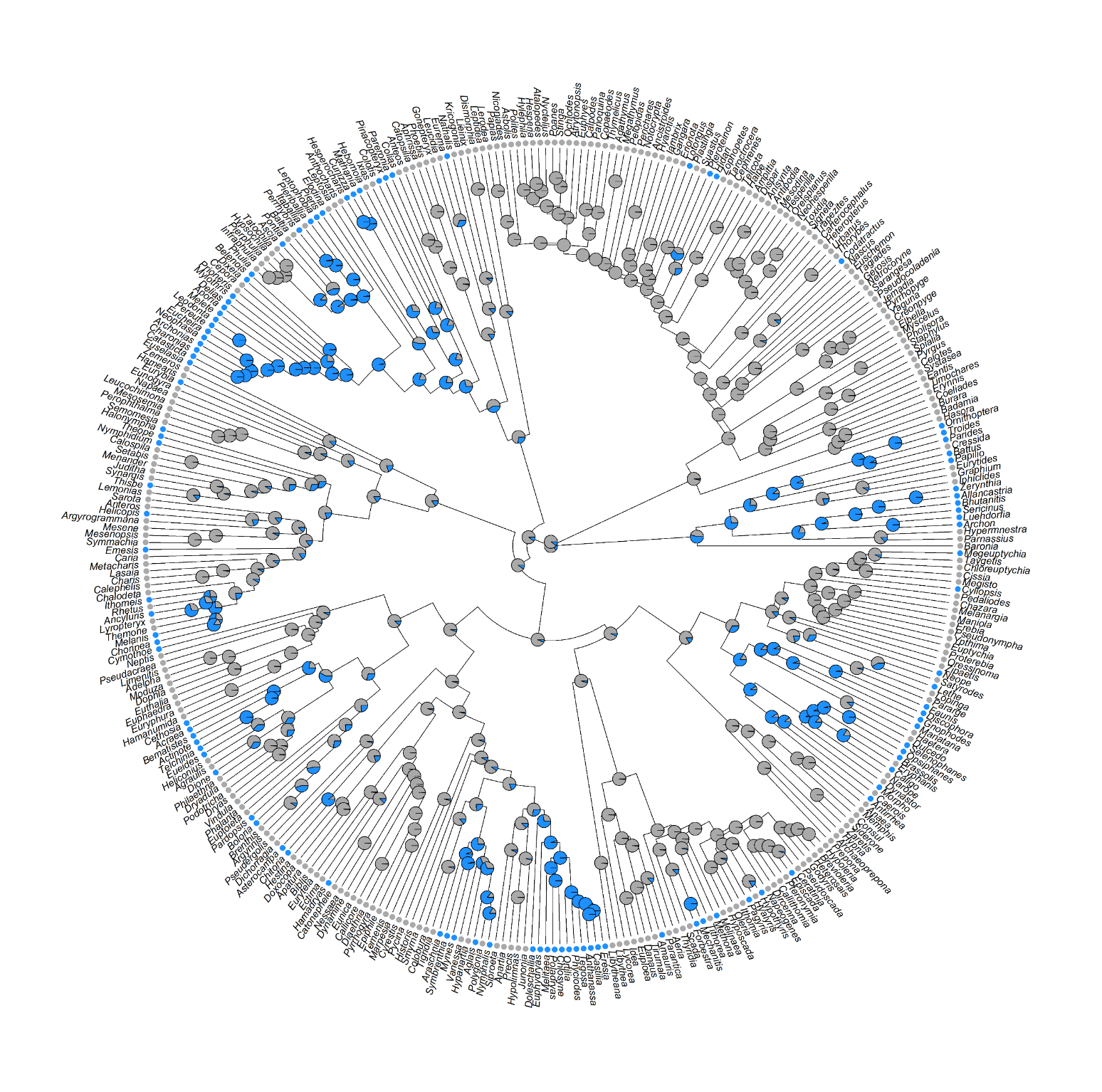

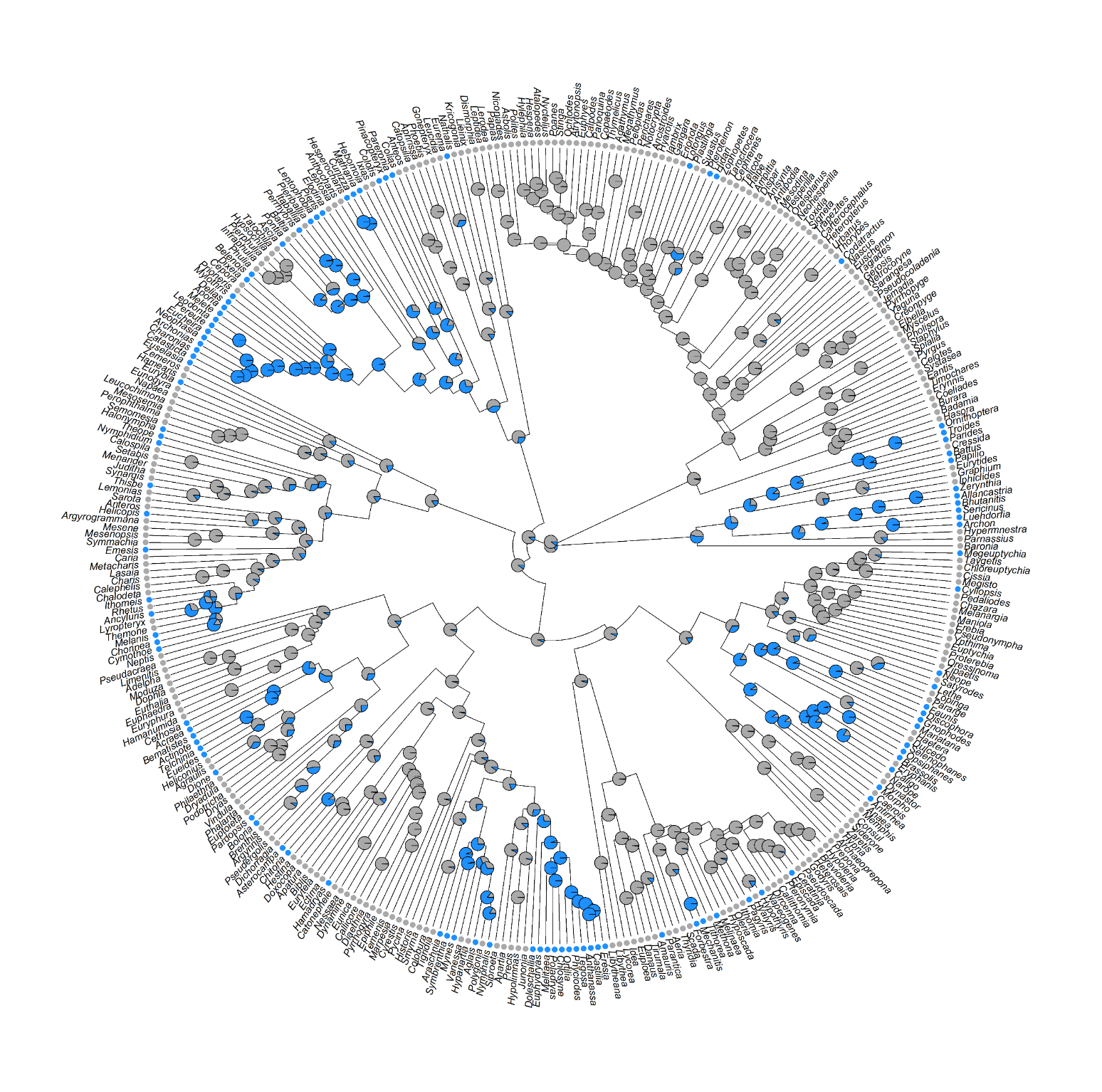

**Hesperiidae**

**Papilionidae**

**Nymphalidae**

**Riodinidae**

**Pieridae**

**Figure S2**

Genus-level butterfly phylogeny adapted from Chazot *et al.* (2019) showing estimated transitions to gregarious behavior (blue nodes and tips) from solitariness (grey nodes and tips). Tips are recorded as gregarious if this behavior is present at any point in larval development. The five separate families are highlighted by color and labelled.

| **Supplementary results tables**  **Table S6.** Rates of transition between larval character states of separate traits across the butterfly phylogeny. Also displayed are the loglikelihoods (LogL) and AIC values of the equal (or null) model (which assumes equal transition rates across the phylogeny) and the All Rates Different (ARD) model (which assumes transition rates vary across the phylogeny). Results from comparison between the two models using a loglikelihood ratio test are given, where a significant difference (p < 0.05) confirms that the ARD model is accepted. | | | | | | |
| --- | --- | --- | --- | --- | --- | --- |
| **Trait** | **Character state** | **Transition rate to other state** | **Model LogL / AIC** | | **LogL ratio** | ***p* value** |
|  |  |  | **Equal** | **ARD** |  |  |
| Social behaviour | Solitary | 0.006 | -177.958 / 357.915 | -174.980 / 353.960 | 5.955 | 0.015 |
|  | Gregarious | 0.020 |  |  |  |  |
| Colour pattern | Cryptic | 0.003 | -136.015 / 274.030 | -130.395 / 264.789 | 11.241 | 0.001 |
|  | Aposematic | 0.016 |  |  |  |  |
| Structural defence | No defence | 0.004 | -84.425 / 170.850 | -84.418 / 172.835 | 0.015 | 0.902 |
|  | Defence | 0.004 |  |  |  |  |
| Host plant | Trees | 0.011 | -152.562 / 307.124 | -152.402 / 308.805 | 0.319 | 0.572 |
|  | Non-trees | 0.011 |  |  |  |  |

| **Table S7.** Coevolutionary rates of transition between character states of larval traits across the butterfly phylogeny. Transitions away from the state in the ‘Character state’ column happen fastest in the presence of the state in the ‘Coevolution state’ column. Also displayed are the loglikelihoods (LogL) and AIC values of the independent (or null) model (which assumes independent evolution of traits) and the dependent model (which assumes the evolution of one trait is influenced by another). Results from comparison between the two models using a loglikelihood ratio test are given, where a significant difference (p < 0.05) confirms that the dependent model is accepted. | | | | | | | |
| --- | --- | --- | --- | --- | --- | --- | --- |
| **Trait** | **Character state** | **Coevolution trait** | **Coevolution state** | **Model LogL / AIC** | | **LogL ratio** | ***p* value** |
|  |  |  |  | **Independent** | **Dependent** |  |  |
| **Social behaviour** | Solitary  Gregarious | Colour pattern | Aposematic  Cryptic | -247.382 / 502.763 | -235.914 / 483.827 | 22.936 | < 0.001 |
|  | Solitary  Gregarious | Structural defence | Defence  No defence | -182.494 / 372.988 | -179.052 / 370.104 | 6.884 | 0.032 |
|  | Solitary  Gregarious | Host plant | Independent | -280.639 / 569.279 | -279.608 / 571.217 | 2.062 | 0.357 |
| **Colour pattern** | Cryptic  Aposematic | Social behaviour | Gregarious  Solitary | -247.382 / 502.763 | -236.218 / 484.436 | 22.327 | < 0.001 |
|  | Cryptic  Aposematic | Structural defence | Defence  No defence | -191.580 / 391.160 | -183.579 / 379.158 | 16.002 | < 0.001 |
|  | Cryptic  Aposematic | Host plant | Non-trees | -235.190 / 478.380 | -224.675 / 461.351 | 21.029 | < 0.001 |
| **Structural defence** | No defence  Defence | Social behaviour | Gregarious  Solitary | -182.494 / 372.988 | -176.659 / 365.317 | 11.671 | 0.003 |
|  | No defence  Defence | Colour pattern | Aposematic  Cryptic | -191.580 / 391.160 | -179.771 / 371.541 | 23.619 | < 0.001 |
|  | No defence  Defence | Host plant | Independent | -185.488 / 378.977 | -185.039 / 382.079 | 0.898 | 0.638 |
| **Host plant** | Trees  Non-trees | Social behaviour | Independent | -280.639 / 569.279 | -280.076 / 572.151 | 1.128 | 0.569 |
|  | Trees  Non-trees | Colour pattern | Independent | -235.190 / 478.380 | -233.298 / 478.597 | 3.783 | 0.151 |
|  | Trees  Non-trees | Structural defence | Independent | -185.488 / 378.977 | -185.075 / 383.150 | 0.827 | 0.661 |

| **Table S8.** The percentage of surviving targets in each treatment across two field experiments. Also displayed are the combined data from both experiments. Asterisks denote targets in a group size level which survived significantly better than those in the other group size. | | | | | |
| --- | --- | --- | --- | --- | --- |
| **Dataset** | **Target color** | **Background color** | **Group size** | | **Test stat / p value** |
|  |  |  | Solitary | Grouped |  |
| Background color | Cryptic | Green | 32% | 37% | Z = -1.250 / *p* = 0.211 |
|  | Cryptic | White | 44% | 38% | Z = -1.607 / *p* = 0.108 |
|  | Aposematic | Green | 45% | 65% * | Z = 3.282 / *p* = 0.001 |
|  | Aposematic | White | 48% | 50% | Z = 0.338 / *p* = 0.736 |
| Body size | Cryptic | Green | 67% * | 40% | Z = -13.240 / *p <* 0.001 |
|  | Aposematic | Green | 68% | 68% * | Z = 2.255 / *p* = 0.024 |
| Combined | Cryptic | Green | 52% * | 40% | Z = -2.725 / *p* = 0.006 |
|  | Aposematic | Green | 54% | 68% * | Z = 2.656 / *p* = 0.008 |
