## Supplemental R scripts for "Warning coloration, body size and the evolution of gregarious behavior in butterfly larvae": All_trees_creation.R

```
rm(list=ls())
library(ape)
library(openxlsx)
setwd('C:\\Users\\cm15245\\OneDrive - University of Bristol\\Documents\\(0) PhD\\Comparative stuff\\Data\\(0) NEW datasets')
phy<-ape::read.nexus(file = 'S_3b_core_analysis_median_ages.TRE')

splitnames <- strsplit(phy$tip.label, "_")
splitnames

#For this to work, in excel doc changed 'Hallonympha' to 'Halonympha', changed 'Psuedonascus' to 'Nascus'
#and removed 'Celaenorrhinus'.

excel <- read.xlsx(xlsxFile = "Tree creation data.xlsx", sheet = 1)
head(excel)
Keep_tips<-excel$phylo
Keep_tips

Keep_tips <- unlist(lapply(splitnames, function(x){
  x[x %in% excel$phylo]
  }))
Keep_tips

excel$phylo %in% Keep_tips

## change the tip labels into same format
excelG <- read.xlsx(xlsxFile = "AllGenera_ordered.xlsx",sheet = 1)
head(excelG)

phy$tip.label<-excelG$Genus

Keep_tips %in% phy$tip.label
phy$tip.label %in% Keep_tips

##
summary(phy)

rmvtips <- c(phy$tip.label[!phy$tip.label %in% Keep_tips])
phy<-drop.tip(phy,rmvtips,trim.internal = TRUE,subtree = FALSE,root.edge = 0,rooted = is.rooted(phy),collapse.singles = TRUE,interactive = FALSE)
summary(phy)

write.nexus(phy,file = 'SB_onlyTree.nex')

##################################################################

# Colour pattern
phy<-ape::read.nexus(file = 'S_3b_core_analysis_median_ages.TRE')

splitnames <- strsplit(phy$tip.label, "_")

excel <- read.xlsx(xlsxFile = "Tree creation data.xlsx", sheet = 2)
head(excel)
Keep_tips<-excel$phylo
Keep_tips

Keep_tips <- unlist(lapply(splitnames, function(x){
  x[x %in% excel$phylo]
}))
Keep_tips

excel$phylo %in% Keep_tips

## change the tip labels into same format
excelG <- read.xlsx(xlsxFile = "AllGenera_ordered.xlsx",sheet = 1)
phy$tip.label<-excelG$Genus

Keep_tips %in% phy$tip.label
phy$tip.label %in% Keep_tips

##
summary(phy)

rmvtips <- c(phy$tip.label[!phy$tip.label %in% Keep_tips])
phy<-drop.tip(phy,rmvtips,trim.internal = TRUE,subtree = FALSE,root.edge = 0,rooted = is.rooted(phy),collapse.singles = TRUE,interactive = FALSE)
summary(phy)

write.nexus(phy,file = 'SB_CPTree.nex')

##################################################################

# Structural defence
phy<-ape::read.nexus(file = 'S_3b_core_analysis_median_ages.TRE')

splitnames <- strsplit(phy$tip.label, "_")

excel <- read.xlsx(xlsxFile = "Tree creation data.xlsx", sheet = 3)
head(excel)
Keep_tips<-excel$phylo
Keep_tips

Keep_tips <- unlist(lapply(splitnames, function(x){
  x[x %in% excel$phylo]
}))
Keep_tips

excel$phylo %in% Keep_tips

## change the tip labels into same format
excelG <- read.xlsx(xlsxFile = "AllGenera_ordered.xlsx",sheet = 1)
phy$tip.label<-excelG$Genus

Keep_tips %in% phy$tip.label
phy$tip.label %in% Keep_tips

##
summary(phy)

rmvtips <- c(phy$tip.label[!phy$tip.label %in% Keep_tips])
phy<-drop.tip(phy,rmvtips,trim.internal = TRUE,subtree = FALSE,root.edge = 0,rooted = is.rooted(phy),collapse.singles = TRUE,interactive = FALSE)
summary(phy)

write.nexus(phy,file = 'SB_SDTree.nex')

##################################################################

# Body size 
phy<-ape::read.nexus(file = 'S_3b_core_analysis_median_ages.TRE')

splitnames <- strsplit(phy$tip.label, "_")

excel <- read.xlsx(xlsxFile = "Tree creation data.xlsx", sheet = 4)
head(excel)
Keep_tips<-excel$phylo
Keep_tips

Keep_tips <- unlist(lapply(splitnames, function(x){
  x[x %in% excel$phylo]
}))
Keep_tips

excel$phylo %in% Keep_tips

## change the tip labels into same format
excelG <- read.xlsx(xlsxFile = "AllGenera_ordered.xlsx",sheet = 1)
phy$tip.label<-excelG$Genus

Keep_tips %in% phy$tip.label
phy$tip.label %in% Keep_tips

##
summary(phy)

rmvtips <- c(phy$tip.label[!phy$tip.label %in% Keep_tips])
phy<-drop.tip(phy,rmvtips,trim.internal = TRUE,subtree = FALSE,root.edge = 0,rooted = is.rooted(phy),collapse.singles = TRUE,interactive = FALSE)
summary(phy)

write.nexus(phy,file = 'SB_BSTree.nex')

##################################################################

# Host plant
phy<-ape::read.nexus(file = 'S_3b_core_analysis_median_ages.TRE')

splitnames <- strsplit(phy$tip.label, "_")

excel <- read.xlsx(xlsxFile = "Tree creation data.xlsx", sheet = 5)
head(excel)
Keep_tips<-excel$phylo
Keep_tips

Keep_tips <- unlist(lapply(splitnames, function(x){
  x[x %in% excel$phylo]
}))
Keep_tips

excel$phylo %in% Keep_tips

## change the tip labels into same format
excelG <- read.xlsx(xlsxFile = "AllGenera_ordered.xlsx",sheet = 1)
phy$tip.label<-excelG$Genus

Keep_tips %in% phy$tip.label
phy$tip.label %in% Keep_tips

##
summary(phy)

rmvtips <- c(phy$tip.label[!phy$tip.label %in% Keep_tips])
phy<-drop.tip(phy,rmvtips,trim.internal = TRUE,subtree = FALSE,root.edge = 0,rooted = is.rooted(phy),collapse.singles = TRUE,interactive = FALSE)
summary(phy)

write.nexus(phy,file = 'SB_HPTree.nex')

##################################################################

# Colour and Structural defence
phy<-ape::read.nexus(file = 'S_3b_core_analysis_median_ages.TRE')

splitnames <- strsplit(phy$tip.label, "_")

excel <- read.xlsx(xlsxFile = "Tree creation data.xlsx", sheet = 6)
head(excel)
Keep_tips<-excel$phylo
Keep_tips

Keep_tips <- unlist(lapply(splitnames, function(x){
  x[x %in% excel$phylo]
}))
Keep_tips

excel$phylo %in% Keep_tips

## change the tip labels into same format
excelG <- read.xlsx(xlsxFile = "AllGenera_ordered.xlsx",sheet = 1)
phy$tip.label<-excelG$Genus

Keep_tips %in% phy$tip.label
phy$tip.label %in% Keep_tips

##
summary(phy)

rmvtips <- c(phy$tip.label[!phy$tip.label %in% Keep_tips])
phy<-drop.tip(phy,rmvtips,trim.internal = TRUE,subtree = FALSE,root.edge = 0,rooted = is.rooted(phy),collapse.singles = TRUE,interactive = FALSE)
summary(phy)

write.nexus(phy,file = 'CP_SDTree.nex')

##################################################################

# Colour and Body size
phy<-ape::read.nexus(file = 'S_3b_core_analysis_median_ages.TRE')

splitnames <- strsplit(phy$tip.label, "_")

excel <- read.xlsx(xlsxFile = "Tree creation data.xlsx", sheet = 7)
head(excel)
Keep_tips<-excel$phylo
Keep_tips

Keep_tips <- unlist(lapply(splitnames, function(x){
  x[x %in% excel$phylo]
}))
Keep_tips

excel$phylo %in% Keep_tips

## change the tip labels into same format
excelG <- read.xlsx(xlsxFile = "AllGenera_ordered.xlsx",sheet = 1)
phy$tip.label<-excelG$Genus

Keep_tips %in% phy$tip.label
phy$tip.label %in% Keep_tips

##
summary(phy)

rmvtips <- c(phy$tip.label[!phy$tip.label %in% Keep_tips])
phy<-drop.tip(phy,rmvtips,trim.internal = TRUE,subtree = FALSE,root.edge = 0,rooted = is.rooted(phy),collapse.singles = TRUE,interactive = FALSE)
summary(phy)

write.nexus(phy,file = 'CP_BSTree.nex')

##################################################################

# Colour and Host plant
phy<-ape::read.nexus(file = 'S_3b_core_analysis_median_ages.TRE')

splitnames <- strsplit(phy$tip.label, "_")

excel <- read.xlsx(xlsxFile = "Tree creation data.xlsx", sheet = 8)
head(excel)
Keep_tips<-excel$phylo
Keep_tips

Keep_tips <- unlist(lapply(splitnames, function(x){
  x[x %in% excel$phylo]
}))
Keep_tips

excel$phylo %in% Keep_tips

## change the tip labels into same format
excelG <- read.xlsx(xlsxFile = "AllGenera_ordered.xlsx",sheet = 1)
phy$tip.label<-excelG$Genus

Keep_tips %in% phy$tip.label
phy$tip.label %in% Keep_tips

##
summary(phy)

rmvtips <- c(phy$tip.label[!phy$tip.label %in% Keep_tips])
phy<-drop.tip(phy,rmvtips,trim.internal = TRUE,subtree = FALSE,root.edge = 0,rooted = is.rooted(phy),collapse.singles = TRUE,interactive = FALSE)
summary(phy)

write.nexus(phy,file = 'CP_HPTree.nex')

##################################################################

# Structural defence and Host plant
phy<-ape::read.nexus(file = 'S_3b_core_analysis_median_ages.TRE')

splitnames <- strsplit(phy$tip.label, "_")

excel <- read.xlsx(xlsxFile = "Tree creation data.xlsx", sheet = 9)
head(excel)
Keep_tips<-excel$phylo
Keep_tips

Keep_tips <- unlist(lapply(splitnames, function(x){
  x[x %in% excel$phylo]
}))
Keep_tips

excel$phylo %in% Keep_tips

## change the tip labels into same format
excelG <- read.xlsx(xlsxFile = "AllGenera_ordered.xlsx",sheet = 1)
phy$tip.label<-excelG$Genus

Keep_tips %in% phy$tip.label
phy$tip.label %in% Keep_tips

##
summary(phy)

rmvtips <- c(phy$tip.label[!phy$tip.label %in% Keep_tips])
phy<-drop.tip(phy,rmvtips,trim.internal = TRUE,subtree = FALSE,root.edge = 0,rooted = is.rooted(phy),collapse.singles = TRUE,interactive = FALSE)
summary(phy)

write.nexus(phy,file = 'SD_HPTree.nex')
##################################################################

# Social, Colour and Body size
phy<-ape::read.nexus(file = 'S_3b_core_analysis_median_ages.TRE')

splitnames <- strsplit(phy$tip.label, "_")

excel <- read.xlsx(xlsxFile = "Tree creation data.xlsx", sheet = 10)
head(excel)
Keep_tips<-excel$phylo
Keep_tips

Keep_tips <- unlist(lapply(splitnames, function(x){
  x[x %in% excel$phylo]
}))
Keep_tips

excel$phylo %in% Keep_tips

## change the tip labels into same format
excelG <- read.xlsx(xlsxFile = "AllGenera_ordered.xlsx",sheet = 1)
phy$tip.label<-excelG$Genus

Keep_tips %in% phy$tip.label
phy$tip.label %in% Keep_tips

##
summary(phy)

rmvtips <- c(phy$tip.label[!phy$tip.label %in% Keep_tips])
phy<-drop.tip(phy,rmvtips,trim.internal = TRUE,subtree = FALSE,root.edge = 0,rooted = is.rooted(phy),collapse.singles = TRUE,interactive = FALSE)
summary(phy)

write.nexus(phy,file = 'SB_CP_BSTree.nex')

##################################################################

# All traits
phy<-ape::read.nexus(file = 'S_3b_core_analysis_median_ages.TRE')

splitnames <- strsplit(phy$tip.label, "_")

excel <- read.xlsx(xlsxFile = "Tree creation data.xlsx", sheet = 11)
head(excel)
Keep_tips<-excel$phylo
Keep_tips

Keep_tips <- unlist(lapply(splitnames, function(x){
  x[x %in% excel$phylo]
}))
Keep_tips

excel$phylo %in% Keep_tips

## change the tip labels into same format
excelG <- read.xlsx(xlsxFile = "AllGenera_ordered.xlsx",sheet = 1)
phy$tip.label<-excelG$Genus

Keep_tips %in% phy$tip.label
phy$tip.label %in% Keep_tips

##
summary(phy)

rmvtips <- c(phy$tip.label[!phy$tip.label %in% Keep_tips])
phy<-drop.tip(phy,rmvtips,trim.internal = TRUE,subtree = FALSE,root.edge = 0,rooted = is.rooted(phy),collapse.singles = TRUE,interactive = FALSE)
summary(phy)

write.nexus(phy,file = 'All_TraitsTree.nex')

##################################################################

# Larval body size and Wingspan
phy<-ape::read.nexus(file = 'S_3b_core_analysis_median_ages.TRE')

splitnames <- strsplit(phy$tip.label, "_")

excel <- read.xlsx(xlsxFile = "Tree creation data.xlsx", sheet = 12)
head(excel)
Keep_tips<-excel$phylo
Keep_tips

Keep_tips <- unlist(lapply(splitnames, function(x){
  x[x %in% excel$phylo]
}))
Keep_tips

excel$phylo %in% Keep_tips

## change the tip labels into same format
excelG <- read.xlsx(xlsxFile = "AllGenera_ordered.xlsx",sheet = 1)
phy$tip.label<-excelG$Genus

Keep_tips %in% phy$tip.label
phy$tip.label %in% Keep_tips

##
summary(phy)

rmvtips <- c(phy$tip.label[!phy$tip.label %in% Keep_tips])
phy<-drop.tip(phy,rmvtips,trim.internal = TRUE,subtree = FALSE,root.edge = 0,rooted = is.rooted(phy),collapse.singles = TRUE,interactive = FALSE)
summary(phy)

write.nexus(phy,file = 'LBS_WSTree.nex')
```
