## Supplemental R scripts for "Warning coloration, body size and the evolution of gregarious behavior in butterfly larvae": Field_data_analyses.R

```
# Background colour experiment

rm(list=ls())

library(openxlsx)
library(lme4)
library(ggplot2)

setwd('C:\\Users\\cm15245\\OneDrive - University of Bristol\\Documents\\(0) PhD\\Bitrex experiments')

data <- read.delim(file = "Experiment_1_background.txt",header = T)

str(data)

data$ID <- factor(data$ID)
data$Block <- factor(data$Block)
data$Background <- factor(data$Background)
data$Colour <- factor(data$Colour)
data$Group.size <- factor(data$Group.size)
data$Survival <- factor(data$Survival)

# Remove NA values from data
data<- na.omit(data)
summary(data$Survival)

str(data)

# Three-way interaction to start
m <- glmer(Survival~Background*Colour*Group.size + (1|ID)+ (1|Block), family = 'binomial', control=glmerControl(optimizer="bobyqa",optCtrl=list(maxfun=2e5)), data = data)
mm <- glmer(Survival~Background*Colour*Group.size + (1|ID), family = 'binomial', control=glmerControl(optimizer="bobyqa",optCtrl=list(maxfun=2e5)), data = data)
anova (m, mm)
# No difference in models, use simpler model mm

# Also try a model with just Block as a random effect:
mmm <- glmer(Survival~Background*Colour*Group.size + (1|Block), family = 'binomial', control=glmerControl(optimizer="bobyqa",optCtrl=list(maxfun=2e5)), data = data)
anova(m, mmm) # Significantly different

m.1<- update(mm,~. -Background:Colour:Group.size)
anova (mm, m.1) 
# The three-way interaction is significant
#Look at individual factors and the interaction between the other two factors.

#------------------------------------
# Colour/palatability - cryptic
m1a <- glmer(Survival~Background*Group.size + (1|ID), family = 'binomial', control=glmerControl(optimizer="bobyqa",optCtrl=list(maxfun=2e5)), data = subset(data, Colour == 'c'))
m1a.1<- update(m1a,~. -Background:Group.size)
anova (m1a, m1a.1) 
# Interaction is not significant
drop1(m1a.1, test='Chisq')# Shows to drop background
m1a.2<- update(m1a.1,~. -Background)
drop1(m1a.2, test='Chisq')
summary(m1a.2)
## Group.size is pretty marginal; not quite significant on the preferred LRT but just sig on the (less preferred) z-test.
## Easier to interpret with tables
round(prop.table(table(subset(data, Colour == 'c' & Group.size == 1)$Survival, subset(data, Colour == 'c' & Group.size == 1)$Background), margin=2), 2)
round(prop.table(table(subset(data, Colour == 'c' & Group.size == 15)$Survival, subset(data, Colour == 'c' & Group.size == 15)$Background), margin=2), 2)
## So, background has no effect on survival for 'c' prey, regardless of group size. About 40% survive in all groups.


# Colour/palatability - aposematic
## Again, try the model with Block and ID first
m1b <- glmer(Survival~Background*Group.size + (1|ID), family = 'binomial', control=glmerControl(optimizer="bobyqa",optCtrl=list(maxfun=2e5)), data = subset(data, Colour == 'a'))
m1b.1<- update(m1b,~. -Background:Group.size)
anova (m1b, m1b.1)
# Interaction is significant, so split

## Group size 1. Can't fit ID as there is only 1 prey.
m1b.1.1 <- glmer(Survival~Background + (1|Block), family = 'binomial', control=glmerControl(optimizer="bobyqa",optCtrl=list(maxfun=2e5)), data = subset(data, Colour == 'a' & Group.size == 1))
drop1(m1b.1.1, test='Chisq')
summary(m1b.1.1)
round(prop.table(table(subset(data, Colour == 'a' & Group.size == 1)$Survival, subset(data, Colour == 'a' & Group.size == 1)$Background), margin=2), 2)
## So, background has no effect on survival for single aposematic prey.

## Group size 15. 
m1b2.1.15 <- glmer(Survival~Background + (1|ID), family = 'binomial', control=glmerControl(optimizer="bobyqa",optCtrl=list(maxfun=2e5)), data = subset(data, Colour == 'a' & Group.size == 15))
drop1(m1b2.1.15, test='Chisq')
summary(m1b2.1.15)
round(prop.table(table(subset(data, Colour == 'a' & Group.size == 15)$Survival, subset(data, Colour == 'a' & Group.size == 15)$Background), margin=2), 2)
## So, g background improves survival for grouped aposematic prey more than w.

#------------------------------------
## Now splitting by group size.
# Group size - solitary (1)
# We can't fit ID for solitary prey
m2a <- glmer(Survival~Colour*Background + (1|Block), family = 'binomial', control=glmerControl(optimizer="bobyqa",optCtrl=list(maxfun=2e5)), data = subset(data, Group.size == '1'))
m2a.1<- update(m2a,~. -Colour:Background)
anova (m2a, m2a.1) 
# Interaction is not significant
drop1(m2a.1, test='Chisq')
m2a.2<- update(m2a.1,~. -Background)
drop1(m2a.2, test='Chisq')
summary(m2a.2)
## For solitary prey, neither colour nor background affect survival.

# Group size - grouped (15)
## We should try and model with both random effects first
m2b <- glmer(Survival~Colour*Background + (1|ID), family = 'binomial', control=glmerControl(optimizer="bobyqa",optCtrl=list(maxfun=2e5)), data = subset(data, Group.size == '15'))
m2b.1<- update(m2b,~. -Colour:Background)
anova (m2b, m2b.1)
# Interaction is significant, so keep it in model and split by colour

# Cryptic
m2b.g <- glmer(Survival~Background + (1|ID), family = 'binomial', control=glmerControl(optimizer="bobyqa",optCtrl=list(maxfun=2e5)), data = subset(data, Group.size == '15' & Colour =='c'))
drop1(m2b.g, test='Chisq')
summary(m2b.g)
# No difference in survival between backgrounds for grouped, cryptic targets.

# Aposematic
m2b.w <- glmer(Survival~Background + (1|ID), family = 'binomial', control=glmerControl(optimizer="bobyqa",optCtrl=list(maxfun=2e5)), data = subset(data, Group.size == '15' & Colour =='a'))
drop1(m2b.w, test='Chisq')
summary(m2b.w)
round(prop.table(table(subset(data, Colour == 'a' & Group.size == 15)$Survival, subset(data, Colour == 'a' & Group.size == 15)$Background), margin=2), 2)
# So, g background improves survival for grouped aposematic prey more than w.

#------------------------------------
# Background - green
m3a <- glmer(Survival~Colour*Group.size + (1|ID), family = 'binomial', control=glmerControl(optimizer="bobyqa",optCtrl=list(maxfun=2e5)), data = subset(data, Background == 'g'))
m3a.1<- update(m3a,~. -Colour:Group.size)
anova (m3a, m3a.1)
# Interaction is significant, so split data

# Split by colour 
# Cryptic
m4a.c <- glmer(Survival~Group.size + (1|ID), family = 'binomial', control=glmerControl(optimizer="bobyqa",optCtrl=list(maxfun=2e5)), data = subset(data, Background == 'g' & Colour == 'c'))
drop1(m4a.c, test='Chisq')
# No effect of group size for cryptic prey on a green background.

# Aposematic
m4a.a <- glmer(Survival~Group.size + (1|ID), family = 'binomial', control=glmerControl(optimizer="bobyqa",optCtrl=list(maxfun=2e5)), data = subset(data, Background == 'g' & Colour == 'a'))
drop1(m4a.a, test='Chisq')
# group size effects survival of aposematic targets on a green background.
round(prop.table(table(subset(data, Background == 'g' & Colour == 'a')$Survival, subset(data, Background == 'g' & Colour == 'a')$Group.size), margin=2), 2)
# Grouping improved aposematic target survival on a green background more than being solitary 


#######################################################################################################
#######################################################################################################

# Body size experiment

rm(list=ls())

library(openxlsx)
library(lme4)
library(ggplot2)
library(dotwhisker)
library(dplyr)
library(sjPlot)
library(sjlabelled)
library(sjmisc)
library(multcomp)
setwd('C:\\Users\\cm15245\\OneDrive - University of Bristol\\Documents\\(0) PhD\\Bitrex experiments')

data <- read.delim(file = "Experiment_2_bodysize.txt",header = T)
str(data)

data$ID <- factor(data$ID)
data$Block <- factor(data$Block)
data$Colour <- factor(data$Colour)
data$Size <- factor(data$Size)
data$Group <- factor(data$Group)
data$Survived <- factor(data$Survived)

# Remove NA values from data
data<- na.omit(data)
summary(data$Survived)

str(data)

# Three-way interaction to start
m <- glmer(Survived~Colour*Group*Size + (1|ID)+ (1|Block), family = 'binomial', control=glmerControl(optimizer="bobyqa",optCtrl=list(maxfun=2e5)), data = data)
mm <- glmer(Survived~Colour*Group*Size + (1|ID), family = 'binomial', control=glmerControl(optimizer="bobyqa",optCtrl=list(maxfun=2e5)), data = data)
anova (m, mm)
# Sig diff between models
m.1<- update(m,~. -Colour:Group:Size)
anova (m, m.1) 
# Three-way interaction is not significant

drop1(m.1, test='Chisq')

m.2<- update(m.1,~. -Colour:Size)
drop1(m.2, test='Chisq')

m.3<- update(m.2,~. -Group:Size)
drop1(m.3, test='Chisq')

m.4<- update(m.3,~. -Size)
drop1(m.4, test='Chisq')
# No effect of size. now split data by colour and check effect of group on each.

# Cryptic
m1 <- glmer(Survived~Group + (1|ID), family = 'binomial', control=glmerControl(optimizer="bobyqa",optCtrl=list(maxfun=2e5)), data = subset(data, Colour == 'c'))
drop1(m1, test='Chisq')
summary(m1) 
round(prop.table(table(subset(data, Colour == 'c')$Survived, subset(data, Colour == 'c')$Group), margin=2), 2)
# Being solitary improves cryptic prey survival far more than grouping.

# Aposematic
m2 <- glmer(Survived~Group + (1|ID), family = 'binomial', control=glmerControl(optimizer="bobyqa",optCtrl=list(maxfun=2e5)), data = subset(data, Colour == 'a'))
drop1(m2, test='Chisq')
summary(m2)
round(prop.table(table(subset(data, Colour == 'a')$Survived, subset(data, Colour == 'a')$Group), margin=2), 2)
# Both groups survived equally well


#######################################################################################################
#######################################################################################################

# Combined data

rm(list=ls())

library(openxlsx)
library(lme4)
library(ggplot2)

setwd('C:\\Users\\cm15245\\OneDrive - University of Bristol\\Documents\\(0) PhD\\Bitrex experiments')

data <- read.delim(file = "Experiment_3_combined.txt",header = T)
str(data)

data$ID <- factor(data$ID)
data$Block <- factor(data$Block)
data$Colour <- factor(data$Colour)
data$Group <- factor(data$Group)
data$Survived <- factor(data$Survived)

# Remove NA values from data
data<- na.omit(data)
summary(data$Survived)

str(data)

# Colour * Group
modd <- glmer(Survived~Colour*Group + (1|ID)+ (1|Block), family = 'binomial', control=glmerControl(optimizer="bobyqa",optCtrl=list(maxfun=2e5)), data = data)
mod <- glmer(Survived~Colour*Group + (1|ID), family = 'binomial', control=glmerControl(optimizer="bobyqa",optCtrl=list(maxfun=2e5)), data = data)
anova (modd, mod)
# No difference between models, so use simpler without Block

mod.1<- update(mod,~. -Colour:Group)
anova (mod, mod.1) 
# Interaction is significant, so split by colour and test effect of group on each

# Cryptic
mod1 <- glmer(Survived~Group + (1|ID), family = 'binomial', control=glmerControl(optimizer="bobyqa",optCtrl=list(maxfun=2e5)), data = subset(data, Colour == 'c'))
drop1(mod1, test='Chisq') 
summary(mod1) 
round(prop.table(table(subset(data, Colour == 'c')$Survived, subset(data, Colour == 'c')$Group), margin=2), 2)
# No difference between solitary cryptic prey, but grouping lessens cryptic targets survival

# Aposematic
mod2 <- glmer(Survived~Group + (1|ID), family = 'binomial', control=glmerControl(optimizer="bobyqa",optCtrl=list(maxfun=2e5)), data = subset(data, Colour == 'a'))
drop1(mod2, test='Chisq')
summary(mod2) 
round(prop.table(table(subset(data, Colour == 'a')$Survived, subset(data, Colour == 'a')$Group), margin=2), 2)
# No difference between solitary aposematic prey, but grouping increases survival

# Solitary comparison
mod3 <- glmer(Survived~Colour + (1|ID), family = 'binomial', control=glmerControl(optimizer="bobyqa",optCtrl=list(maxfun=2e5)), data = subset(data, Group == '1'))
drop1(mod3, test='Chisq')
## There is a warning "Model is nearly unidentifiable: large eigenvalue ratio"
## This is because, with solitary prey, the prey is only either eaten or not,
## so the random effect of ID has very little variance

## We can compare the variance explained by the mixed model and model without the random effect of ID.
mod3a <- glm(Survived~Colour, family = 'binomial', data = subset(data, Group == '1'))
anova(mod3, mod3a, test='Chisq')

## No difference at all. However, the survival of prey within the same block could still be more similar than you'd expect by chance:
mod3b <- glmer(Survived~Colour + (1|Block), family = 'binomial', control=glmerControl(optimizer="bobyqa",optCtrl=list(maxfun=2e5)), data = subset(data, Group == '1'))
anova(mod3a, mod3b, test='Chisq')
# Again no difference, so can use glm model
drop1(mod3a, test='Chisq')

# Grouped comparison
mod4 <- glmer(Survived~Colour + (1|ID), family = 'binomial', control=glmerControl(optimizer="bobyqa",optCtrl=list(maxfun=2e5)), data = subset(data, Group == '15'))
drop1(mod4, test='Chisq')
summary(mod4)
```
