## Supplemental R scripts for "Warning coloration, body size and the evolution of gregarious behavior in butterfly larvae": MCMCglmms.R

```
rm(list=ls())

library(ape)
library(MCMCglmm)
library(openxlsx)
library(phytools)
setwd('C:\\Users\\cm15245\\OneDrive - University of Bristol\\Documents\\(0) PhD\\Comparative stuff\\Data\\(0) NEW datasets')
phylo<-ape::read.nexus(file = 'All_TraitsTree.nex')
excel <- read.xlsx(xlsxFile = "AllTraits_Bin.xlsx")

excel$SB<- as.factor(excel$SB)
excel$CP<- as.factor(excel$CP)
excel$SD<- as.factor(excel$SD)
excel$HP<- as.factor(excel$HP)
str(excel)
inv.phylo<-inverseA(phylo,nodes="TIPS",scale=TRUE) 
prior<-list(G=list(G1=list(V=1,nu=1,alpha.mu=0,alpha.V=1000)),R=list(V=1,nu=0.002, fix=1))

All_Bin<- MCMCglmm(SB~CP*SD*HP*BS,random=~phylo,family=c("categorical"),ginverse=list(phylo=inv.phylo$Ainv),prior=prior,data=excel,nitt=5100000,burnin=100000,thin=500) 
summary(All_Bin)
windows()
plot(All_Bin) 
prop.table(table(All_Bin$Sol[,2]>0))
#FALSE   TRUE 
#0.9637 0.0363 
save(All_Bin, file = "All_Bin.RData")
load("All_Bin.RData")
lambda <- All_Bin$VCV[,'phylo']/(All_Bin$VCV[,'phylo']+All_Bin$VCV[,'units']) 
mean(lambda)
#0.9999771

# Remove SD
All_Bin1<- MCMCglmm(SB~CP*BS*HP+CP:SD:HP:BS+SD:HP:BS+CP:SD:BS+CP:SD:HP+
                      SD:BS+SD:HP+CP:SD,random=~phylo,family=c("categorical"),ginverse=list(phylo=inv.phylo$Ainv),prior=prior,data=excel,nitt=5100000,burnin=100000,thin=500) 
summary(All_Bin1)
save(All_Bin1, file = "All_Bin1.RData")

# Remove HP : SD
All_Bin2<- MCMCglmm(SB~CP*BS*HP+CP:SD:HP:BS+SD:HP:BS+CP:SD:BS+CP:SD:HP+
                      SD:BS+CP:SD,random=~phylo,family=c("categorical"),ginverse=list(phylo=inv.phylo$Ainv),prior=prior,data=excel,nitt=5100000,burnin=100000,thin=500) 
summary(All_Bin2)
save(All_Bin2, file = "All_Bin2.RData")

# Remove BS
All_Bin3<- MCMCglmm(SB~CP*HP+CP:SD:HP:BS+SD:HP:BS+CP:SD:BS+CP:SD:HP+CP:BS:HP+
                      SD:BS+BS:HP+CP:BS+CP:SD,random=~phylo,family=c("categorical"),ginverse=list(phylo=inv.phylo$Ainv),prior=prior,data=excel,nitt=5100000,burnin=100000,thin=500) 
summary(All_Bin3)
save(All_Bin3, file = "All_Bin3.RData")

# Remove SD : BS
All_Bin4<- MCMCglmm(SB~CP*HP+CP:SD:HP:BS+SD:HP:BS+CP:SD:BS+CP:SD:HP+CP:BS:HP+
                      BS:HP+CP:BS+CP:SD,random=~phylo,family=c("categorical"),ginverse=list(phylo=inv.phylo$Ainv),prior=prior,data=excel,nitt=5100000,burnin=100000,thin=500) 
summary(All_Bin4)
save(All_Bin4, file = "All_Bin4.RData")

# Remove HP : BS
All_Bin5<- MCMCglmm(SB~CP*HP+CP:SD:HP:BS+SD:HP:BS+CP:SD:BS+CP:SD:HP+CP:BS:HP+
                      CP:BS+CP:SD,random=~phylo,family=c("categorical"),ginverse=list(phylo=inv.phylo$Ainv),prior=prior,data=excel,nitt=5100000,burnin=100000,thin=500) 
summary(All_Bin5)
save(All_Bin5, file = "All_Bin5.RData")

# Remove CP : SD
All_Bin6<- MCMCglmm(SB~CP*HP+CP:SD:HP:BS+SD:HP:BS+CP:SD:BS+CP:SD:HP+CP:BS:HP+
                      CP:BS,random=~phylo,family=c("categorical"),ginverse=list(phylo=inv.phylo$Ainv),prior=prior,data=excel,nitt=5100000,burnin=100000,thin=500) 
summary(All_Bin6)
save(All_Bin6, file = "All_Bin6.RData")

# Remove CP : HP : SD
All_Bin7<- MCMCglmm(SB~CP*HP+CP:SD:HP:BS+SD:HP:BS+CP:SD:BS+CP:BS:HP+
                      CP:BS,random=~phylo,family=c("categorical"),ginverse=list(phylo=inv.phylo$Ainv),prior=prior,data=excel,nitt=5100000,burnin=100000,thin=500) 
summary(All_Bin7)
save(All_Bin7, file = "All_Bin7.RData")

# Remove CP : SD : BS : HP
All_Bin8<- MCMCglmm(SB~CP*HP+SD:HP:BS+CP:SD:BS+CP:BS:HP+
                      CP:BS,random=~phylo,family=c("categorical"),ginverse=list(phylo=inv.phylo$Ainv),prior=prior,data=excel,nitt=5100000,burnin=100000,thin=500) 
summary(All_Bin8)
save(All_Bin8, file = "All_Bin8.RData")

# Remove CP
All_Bin9<- MCMCglmm(SB~SD:HP:BS+CP:SD:BS+CP:BS:HP+
                      CP:BS+CP:HP+HP,random=~phylo,family=c("categorical"),ginverse=list(phylo=inv.phylo$Ainv),prior=prior,data=excel,nitt=5100000,burnin=100000,thin=500) 
summary(All_Bin9)
save(All_Bin9, file = "All_Bin9.RData")

# Remove CP : SD : BS
All_Bin10<- MCMCglmm(SB~SD:HP:BS+CP:BS:HP+
                       CP:BS+CP:HP+HP,random=~phylo,family=c("categorical"),ginverse=list(phylo=inv.phylo$Ainv),prior=prior,data=excel,nitt=5100000,burnin=100000,thin=500) 
summary(All_Bin10)
save(All_Bin10, file = "All_Bin10.RData")

# Remove CP : HP
All_Bin11<- MCMCglmm(SB~SD:HP:BS+CP:BS:HP+
                       CP:BS+HP,random=~phylo,family=c("categorical"),ginverse=list(phylo=inv.phylo$Ainv),prior=prior,data=excel,nitt=5100000,burnin=100000,thin=500) 
summary(All_Bin11)
save(All_Bin11, file = "All_Bin11.RData")

# Remove CP : BS : HP
All_Bin12<- MCMCglmm(SB~SD:HP:BS+
                       CP:BS+HP,random=~phylo,family=c("categorical"),ginverse=list(phylo=inv.phylo$Ainv),prior=prior,data=excel,nitt=5100000,burnin=100000,thin=500) 
summary(All_Bin12)
save(All_Bin12, file = "All_Bin12.RData")

# Remove SD : BS : HP
All_Bin13<- MCMCglmm(SB~CP:BS+HP,random=~phylo,family=c("categorical"),ginverse=list(phylo=inv.phylo$Ainv),prior=prior,data=excel,nitt=5100000,burnin=100000,thin=500) 
summary(All_Bin13)
save(All_Bin13, file = "All_Bin13.RData")

# Remove CP : BS
All_Bin14<- MCMCglmm(SB~HP,random=~phylo,family=c("categorical"),ginverse=list(phylo=inv.phylo$Ainv),prior=prior,data=excel,nitt=5100000,burnin=100000,thin=500) 
summary(All_Bin14)
save(All_Bin14, file = "All_Bin14.RData")
# No significant interactions
#             post.mean l-95% CI u-95% CI eff.samp pMCMC
#(Intercept)   -24.206  -97.758   27.008    188.5 0.287
#HP1            -8.433  -36.785   15.292    163.9 0.399


###########################################################################################

# Social behaviour ~ Colour pattern - simple, large model

rm(list=ls())
library(ape)
library(MCMCglmm)
library(openxlsx)
library(phytools)

setwd('C:\\Users\\cm15245\\OneDrive - University of Bristol\\Documents\\(0) PhD\\Comparative stuff\\Data\\(0) NEW datasets')
phylo<-ape::read.nexus(file = 'SB_CPTree.nex')
excel <- read.xlsx(xlsxFile = "SB_CP.xlsx")
excel$SB<- as.factor(excel$SB)
excel$CP<- as.factor(excel$CP)
str(excel)

inv.phylo<-inverseA(phylo,nodes="TIPS",scale=TRUE) 
prior<-list(G=list(G1=list(V=1,nu=1,alpha.mu=0,alpha.V=1000)),R=list(V=1,nu=0.002, fix=1)) 

SB_CP<- MCMCglmm(SB~CP,random=~phylo,family=c("categorical"),ginverse=list(phylo=inv.phylo$Ainv),prior=prior,data=excel,nitt=5100000,burnin=100000,thin=500) 
summary(SB_CP) 
#             post.mean l-95% CI u-95% CI eff.samp  pMCMC  
#(Intercept)   -91.734 -183.062   -7.337    36.23 0.0112 * 
#CP1            35.683    2.166   71.316    85.12 0.0088 **
plot(SB_CP) 
prop.table(table(SB_CP$Sol[,2]>0))
#FALSE   TRUE 
#0.0044 0.9956
save(SB_CP, file = "SB_CP.RData")
load("SB_CP.RData")
lambda <- SB_CP$VCV[,'phylo']/(SB_CP$VCV[,'phylo']+SB_CP$VCV[,'units']) 
mean(lambda)
#0.9996552


###########################################################################################

# Social behaviour ~ Structural defence

rm(list=ls())
library(ape)
library(MCMCglmm)
library(openxlsx)
library(phytools)

setwd('C:\\Users\\cm15245\\OneDrive - University of Bristol\\Documents\\(0) PhD\\Comparative stuff\\Data\\(0) NEW datasets')
phylo<-ape::read.nexus(file = 'SB_SDTree.nex')
excel <- read.xlsx(xlsxFile = "SB_SD.xlsx") 
excel$SB<- as.factor(excel$SB)
excel$SD<- as.factor(excel$SD)
str(excel)

inv.phylo<-inverseA(phylo,nodes="TIPS",scale=TRUE) 
prior<-list(G=list(G1=list(V=1,nu=1,alpha.mu=0,alpha.V=1000)),R=list(V=1,nu=0.002, fix=1)) 

SB_SD<- MCMCglmm(SB~SD,random=~phylo,family=c("categorical"),ginverse=list(phylo=inv.phylo$Ainv),prior=prior,data=excel,nitt=5100000,burnin=100000,thin=500) 
summary(SB_SD) 
#              post.mean l-95% CI u-95% CI eff.samp pMCMC
#(Intercept)  -114.925 -228.304   -5.541    38.08 0.0150 *
#SD1            56.785   -4.716  127.235    46.15 0.0532 .
windows()
plot(SB_SD) 
prop.table(table(SB_SD$Sol[,2]>0))
#FALSE   TRUE
#0.0266 0.9734
save(SB_SD, file = "SB_SD.RData")
load("SB_SD.RData")
lambda <- SB_SD$VCV[,'phylo']/(SB_SD$VCV[,'phylo']+SB_SD$VCV[,'units']) 
mean(lambda)
#0.9998546


###########################################################################################

# Colour pattern ~ Structural defence

rm(list=ls())
library(ape)
library(MCMCglmm)
library(openxlsx)
library(phytools)
setwd('C:\\Users\\cm15245\\OneDrive - University of Bristol\\Documents\\(0) PhD\\Comparative stuff\\Data\\(0) NEW datasets')
phylo<-ape::read.nexus(file = 'CP_SDTree.nex')
excel <- read.xlsx(xlsxFile = "CP_SD.xlsx")

excel$CP<- as.factor(excel$CP)
excel$SD<- as.factor(excel$SD)

str(excel)

inv.phylo<-inverseA(phylo,nodes="TIPS",scale=TRUE) 
prior<-list(G=list(G1=list(V=1,nu=1,alpha.mu=0,alpha.V=1000)),R=list(V=1,nu=0.002, fix=1)) 

CP_SD<- MCMCglmm(CP~SD,random=~phylo,family=c("categorical"),ginverse=list(phylo=inv.phylo$Ainv),prior=prior,data=excel,nitt=5100000,burnin=100000,thin=500) 
summary(CP_SD) 
#               post.mean l-95% CI u-95% CI eff.samp  pMCMC    
#(Intercept)   -90.579 -267.129    2.833    7.227 0.0274 *  
#SD1            95.983    5.589  236.509    4.901 <1e-04 ***
windows()
plot(CP_SD) 
prop.table(table(CP_SD$Sol[,2]>0))
#TRUE 
#1
save(CP_SD, file = "CP_SD.RData")
load("CP_SD.RData")
lambda <- CP_SD$VCV[,'phylo']/(CP_SD$VCV[,'phylo']+CP_SD$VCV[,'units']) 
mean(lambda)
#0.9994211


###########################################################################################

# Social behaviour ~ Colour pattern * Structural defence

rm(list=ls())
library(ape)
library(MCMCglmm)
library(openxlsx)
library(phytools)
setwd('C:\\Users\\cm15245\\OneDrive - University of Bristol\\Documents\\(0) PhD\\Comparative stuff\\Data\\(0) NEW datasets')
phylo<-ape::read.nexus(file = 'CP_SDTree.nex')
excel <- read.xlsx(xlsxFile = "SB_CP_SD.xlsx") 

excel$SB<- as.factor(excel$SB)
excel$CP<- as.factor(excel$CP)
excel$SD<- as.factor(excel$SD)

str(excel)

inv.phylo<-inverseA(phylo,nodes="TIPS",scale=TRUE) 
prior<-list(G=list(G1=list(V=1,nu=1,alpha.mu=0,alpha.V=1000)),R=list(V=1,nu=0.002, fix=1)) 

SB_CPSD<- MCMCglmm(SB~CP*SD,random=~phylo,family=c("categorical"),ginverse=list(phylo=inv.phylo$Ainv),prior=prior,data=excel,nitt=5100000,burnin=100000,thin=500) 
summary(SB_CPSD)
#CP1:SD1        -10.98  -152.95   118.39   102.23 0.8668 
windows()
plot(SB_CPSD) 
prop.table(table(SB_CPSD$Sol[,2]>0))
#FALSE   TRUE
#0.1807 0.8193
save(SB_CPSD, file = "SB_CPSD.RData")
load("SB_CPSD.RData")
lambda <- SB_CPSD$VCV[,'phylo']/(SB_CPSD$VCV[,'phylo']+SB_CPSD$VCV[,'units']) 
mean(lambda)
#0.9999683


###########################################################################################

# Social behaviour ~ Host plant (bin)

rm(list=ls())
library(ape)
library(MCMCglmm)
library(openxlsx)
library(phytools)
setwd('C:\\Users\\cm15245\\OneDrive - University of Bristol\\Documents\\(0) PhD\\Comparative stuff\\Data\\(0) NEW datasets')
phylo<-ape::read.nexus(file = 'SB_HPTree.nex')
excel <- read.xlsx(xlsxFile = "SB_HP.xlsx")

excel$SB<- as.factor(excel$SB)
excel$HP<- as.factor(excel$HP)
str(excel)

inv.phylo<-inverseA(phylo,nodes="TIPS",scale=TRUE) 
prior<-list(G=list(G1=list(V=1,nu=1,alpha.mu=0,alpha.V=1000)),R=list(V=1,nu=0.002, fix=1)) 

SB_HPBin<- MCMCglmm(SB~HP,random=~phylo,family=c("categorical"),ginverse=list(phylo=inv.phylo$Ainv),prior=prior,data=excel,nitt=5100000,burnin=100000,thin=500) 
summary(SB_HPBin)
# HP1           -16.893  -37.780    2.234    265.0 0.068 .
plot(SB_HPBin) 
prop.table(table(SB_HPBin$Sol[,2]>0))
#FALSE   TRUE 
#0.966 0.034
windows()
save(SB_HPBin, file = "SB_HPBin.RData")
load("SB_HPBin.RData")
lambda <- SB_HPBin$VCV[,'phylo']/(SB_HPBin$VCV[,'phylo']+SB_HPBin$VCV[,'units']) 
mean(lambda)
#0.9997906


###########################################################################################

# Social behaviour ~ Host plant (cat)

rm(list=ls())
library(ape)
library(MCMCglmm)
library(openxlsx)
library(phytools)
setwd('C:\\Users\\cm15245\\OneDrive - University of Bristol\\Documents\\(0) PhD\\Comparative stuff\\Data\\(0) NEW datasets')
phylo<-ape::read.nexus(file = 'SB_HPTree.nex')
excel <- read.xlsx(xlsxFile = "SB_HP_Cat.xlsx")

excel$SB<- as.factor(excel$SB)
str(excel)

inv.phylo<-inverseA(phylo,nodes="TIPS",scale=TRUE) 
prior<-list(G=list(G1=list(V=1,nu=1,alpha.mu=0,alpha.V=1000)),R=list(V=1,nu=0.002, fix=1)) 

SB_HPCat<- MCMCglmm(SB~HP,random=~phylo,family=c("categorical"),ginverse=list(phylo=inv.phylo$Ainv),prior=prior,data=excel,nitt=5100000,burnin=100000,thin=500) 
summary(SB_HPCat)
# None significant
plot(SB_HPCat) 
prop.table(table(SB_HPCat$Sol[,2]>0))
#FALSE   TRUE 
#0.8594 0.1406 
windows()
save(SB_HPCat, file = "SB_HPCat.RData")
load("SB_HPCat.RData")
lambda <- SB_HPCat$VCV[,'phylo']/(SB_HPCat$VCV[,'phylo']+SB_HPCat$VCV[,'units']) 
mean(lambda)
#0.9999297


###########################################################################################

# Colour pattern ~ Host plant (cat)

rm(list=ls())
library(ape)
library(MCMCglmm)
library(openxlsx)
library(phytools)

setwd('C:\\Users\\cm15245\\OneDrive - University of Bristol\\Documents\\(0) PhD\\Comparative stuff\\Data\\(0) NEW datasets')
phylo<-ape::read.nexus(file = 'CP_HPTree.nex')
excel <- read.xlsx(xlsxFile = "CP_HP_Cat.xlsx")
excel$CP<- as.factor(excel$CP)
str(excel)

inv.phylo<-inverseA(phylo,nodes="TIPS",scale=TRUE) 
prior<-list(G=list(G1=list(V=1,nu=1,alpha.mu=0,alpha.V=1000)),R=list(V=1,nu=0.002, fix=1)) 

CP_HP<- MCMCglmm(CP~HP,random=~phylo,family=c("categorical"),ginverse=list(phylo=inv.phylo$Ainv),prior=prior,data=excel,nitt=5100000,burnin=100000,thin=500) 
summary(CP_HP) 
windows()
plot(CP_HP) 
prop.table(table(CP_HP$Sol[,2]>0))
#FALSE   TRUE 
#0.9334 0.0666 
save(CP_HP, file = "CP_HP.RData")
load("CP_HP.RData")
lambda <- CP_HP$VCV[,'phylo']/(CP_HP$VCV[,'phylo']+CP_HP$VCV[,'units']) 
mean(lambda)
#0.9998959


###########################################################################################

# Social behaviour ~ Body Size

rm(list=ls())
library(ape)
library(MCMCglmm)
library(openxlsx)
library(phytools)

setwd('C:\\Users\\cm15245\\OneDrive - University of Bristol\\Documents\\(0) PhD\\Comparative stuff\\Data\\(0) NEW datasets')
phylo<-ape::read.nexus(file = 'SB_BSTree.nex')
excel <- read.xlsx(xlsxFile = "SB_BS.xlsx")
excel$SB<- as.factor(excel$SB)
str(excel)

inv.phylo<-inverseA(phylo,nodes="TIPS",scale=TRUE) 
prior<-list(G=list(G1=list(V=1,nu=1,alpha.mu=0,alpha.V=1000)),R=list(V=1,nu=0.002, fix=1)) 

SB_BS<- MCMCglmm(SB~BS,random=~phylo,family=c("categorical"),ginverse=list(phylo=inv.phylo$Ainv),prior=prior,data=excel,nitt=5100000,burnin=100000,thin=500) 
summary(SB_BS) 
#BS        80.77  -55.35   273.09   54.36 0.2208 
windows()
plot(SB_BS) 
prop.table(table(SB_BS$Sol[,2]>0))
# FALSE   TRUE 
#0.1104 0.8896    
save(SB_BS, file = "SB_BS.RData")
load("SB_BS.RData")
lambda <- SB_BS$VCV[,'phylo']/(SB_BS$VCV[,'phylo']+SB_BS$VCV[,'units']) 
mean(lambda)
#0.9998837

# with 'Genus' as random variable instead of phylo:
load("SB_BSTest.RData")
summary(SB_BSTest) # For some reason it's different, still non-sig though

###########################################################################################

# Colour pattern ~ Body Size

rm(list=ls())
library(ape)
library(MCMCglmm)
library(openxlsx)
library(phytools)

setwd('C:\\Users\\cm15245\\OneDrive - University of Bristol\\Documents\\(0) PhD\\Comparative stuff\\Data\\(0) NEW datasets')
phylo<-ape::read.nexus(file = 'CP_BSTree.nex')
excel <- read.xlsx(xlsxFile = "CP_BS.xlsx")
excel$CP<- as.factor(excel$CP)
str(excel)

inv.phylo<-inverseA(phylo,nodes="TIPS",scale=TRUE) 
prior<-list(G=list(G1=list(V=1,nu=1,alpha.mu=0,alpha.V=1000)),R=list(V=1,nu=0.002, fix=1)) 

CP_BS<- MCMCglmm(CP~BS,random=~phylo,family=c("categorical"),ginverse=list(phylo=inv.phylo$Ainv),prior=prior,data=excel,nitt=5100000,burnin=100000,thin=500) 
summary(CP_BS) 
#BS             131.83    15.05   251.76    28.86 8e-04 ***
windows()
plot(CP_BS) 
prop.table(table(CP_BS$Sol[,2]>0))
#FALSE   TRUE 
#0.0004 0.9996
save(CP_BS, file = "CP_BS.RData")
load("CP_BS.RData")
lambda <- CP_BS$VCV[,'phylo']/(CP_BS$VCV[,'phylo']+CP_BS$VCV[,'units']) 
mean(lambda)
#0.9996167


###########################################################################################

# Social behaviour ~ Colour pattern * Body size

rm(list=ls())
library(ape)
library(MCMCglmm)
library(openxlsx)
library(phytools)

setwd('C:\\Users\\cm15245\\OneDrive - University of Bristol\\Documents\\(0) PhD\\Comparative stuff\\Data\\(0) NEW datasets')
phylo<-ape::read.nexus(file = 'SB_CP_BSTree.nex')
excel <- read.xlsx(xlsxFile = "SB_CP_BS.xlsx")
excel$SB<- as.factor(excel$SB)
excel$CP<- as.factor(excel$CP)
str(excel)

inv.phylo<-inverseA(phylo,nodes="TIPS",scale=TRUE) 
prior<-list(G=list(G1=list(V=1,nu=1,alpha.mu=0,alpha.V=1000)),R=list(V=1,nu=0.002, fix=1)) 

SB_CPBS<- MCMCglmm(SB~CP*BS,random=~phylo,family=c("categorical"),ginverse=list(phylo=inv.phylo$Ainv),prior=prior,data=excel,nitt=5100000,burnin=100000,thin=500) 
summary(SB_CPBS) 
#            post.mean l-95% CI u-95% CI eff.samp pMCMC  
#(Intercept)   -338.49  -841.93    32.22    26.19 0.044 *
#CP1            188.95  -270.46   750.79    64.49 0.357  
#BS             128.08   -79.82   392.57    35.57 0.180  
#CP1:BS         -89.43  -413.55   163.56    66.67 0.445
windows()
plot(SB_CPBS) 
prop.table(table(SB_CPBS$Sol[,2]>0))
# FALSE   TRUE 
#0.1785 0.8215   
save(SB_CPBS, file = "SB_CPBS.RData")
load("SB_CPBS.RData")
lambda <- SB_CPBS$VCV[,'phylo']/(SB_CPBS$VCV[,'phylo']+SB_CPBS$VCV[,'units']) 
mean(lambda)
#0.9998125


###########################################################################################

# Solitary ~ Colpattern*Body size
rm(list=ls())
library(ape)
library(MCMCglmm)
library(openxlsx)
library(phytools)
setwd('C:\\Users\\cm15245\\OneDrive - University of Bristol\\Documents\\(0) PhD\\Comparative stuff\\Data\\(0) NEW datasets')
phylo<-ape::read.nexus(file = 'SB_CP_BSTree.nex')
excel <- read.xlsx(xlsxFile = "SB_CP_BS.xlsx")
excel <- subset(excel, excel$SB =='0')

excel$SB<- as.factor(excel$SB)
excel$CP<- as.factor(excel$CP)
str(excel)

inv.phylo<-inverseA(phylo,nodes="TIPS",scale=TRUE) 
prior<-list(G=list(G1=list(V=1,nu=1,alpha.mu=0,alpha.V=1000)),R=list(V=1,nu=0.002, fix=1)) 

Sol_CPBS<- MCMCglmm(CP~BS,random=~phylo,family=c("categorical"),ginverse=list(phylo=inv.phylo$Ainv),prior=prior,data=excel,nitt=5100000,burnin=100000,thin=500) 
summary(Sol_CPBS) 
#BS             211.26    22.91   474.41    12.96 <1e-04 ***
windows()
plot(Sol_CPBS)
prop.table(table(Sol_CPBS$Sol[,2]>0))
#TRUE 
#1
save(Sol_CPBS, file = "Sol_CPBS.RData")
load("Sol_CPBS.RData")
lambda <- Sol_CPBS$VCV[,'phylo']/(Sol_CPBS$VCV[,'phylo']+Sol_CPBS$VCV[,'units']) 
mean(lambda)
#0.9996916


###########################################################################################

# Gregarious ~ Colpattern*Body size
rm(list=ls())
library(ape)
library(MCMCglmm)
library(openxlsx)
library(phytools)
setwd('C:\\Users\\cm15245\\OneDrive - University of Bristol\\Documents\\(0) PhD\\Comparative stuff\\Data\\(0) NEW datasets')
phylo<-ape::read.nexus(file = 'SB_CP_BSTree.nex')
excel <- read.xlsx(xlsxFile = "SB_CP_BS.xlsx")
excel <- subset(excel, excel$SB =='1')

excel$SB<- as.factor(excel$SB)
excel$CP<- as.factor(excel$CP)
str(excel)

inv.phylo<-inverseA(phylo,nodes="TIPS",scale=TRUE) 
prior<-list(G=list(G1=list(V=1,nu=1,alpha.mu=0,alpha.V=1000)),R=list(V=1,nu=0.002, fix=1)) 

Grg_CPBS<- MCMCglmm(CP~BS,random=~phylo,family=c("categorical"),ginverse=list(phylo=inv.phylo$Ainv),prior=prior,data=excel,nitt=5100000,burnin=100000,thin=500) 
summary(Grg_CPBS) 
#BS              99.75  -147.20   430.47    98.25 0.361
windows()
plot(Grg_CPBS)
prop.table(table(Grg_CPBS$Sol[,2]>0))
#FALSE   TRUE 
#0.1806 0.8194 
save(Grg_CPBS, file = "Grg_CPBS.RData")
load("Grg_CPBS.RData")
lambda <- Grg_CPBS$VCV[,'phylo']/(Grg_CPBS$VCV[,'phylo']+Grg_CPBS$VCV[,'units']) 
mean(lambda)
#0.9978235
```
