## Supplemental R scripts for "Warning coloration, body size and the evolution of gregarious behavior in butterfly larvae": PPA.R

```
# All traits (N = 176)

rm(list=ls())
library(phylopath)
library(ape)
require(geiger)
library(phytools)
setwd('C:\\Users\\cm15245\\OneDrive - University of Bristol\\Documents\\(0) PhD\\Comparative stuff\\Data\\(0) NEW datasets')

phy<-ape::read.nexus(file = 'All_TraitsTree.nex')

library(openxlsx)
excel <- read.xlsx(xlsxFile = "AllTraits_Bin.xlsx")
head(excel)
excel$SB<-as.factor(excel$SB)
excel$CP<-as.factor(excel$CP)
excel$SD<-as.factor(excel$SD)
excel$HP<-as.factor(excel$HP)

SB <- excel$SB
names(SB) <- excel$phylo
CP <- excel$CP
names(CP) <- excel$phylo
SD <- excel$SD
names(SD) <- excel$phylo
HP<-excel$HP
names(HP)<-excel$phylo
BL<-excel$BS
names(BL)<-excel$phylo

data<-data.frame(SB,CP,SD,HP,BL)

str(data)

m <- define_model_set(
  null = c(),
  A = c(CP~SD, SB~CP+BL+HP),
  B = c(SD~CP, SB~CP+BL+HP),
  C = c(CP~SD, CP~SB+BL+HP),
  D = c(SD~CP, CP~SB+BL+HP),
  E = c(CP~SD+BL, SB~CP, BL~HP),
  F = c(CP~SD, SB~CP, BL~CP+HP),
  G = c(CP~SD, SB~CP+HP, BL~CP+HP),
  H = c(CP~SD, SB~CP+HP, BL~CP),
  I = c(CP~SD, SB~CP+SD, BL~CP+HP),
  J = c(CP~SD, SB~CP, BL~CP+HP+SB),
  K = c(CP~SD, SB~CP+BL, BL~CP+HP))  

P<-phylo_path(model_set = m, data = data, tree = phy)
P
plot_model_set(m)
s <- summary(P)
s
plot(s)

plot(m$I)
plot(m$F)
plot(m$K)
plot(m$J)
plot(m$G)


b <- best(P) 
plot(b)
b_ci <- best(P, boot = 500)
b_ci
plot(b_ci)

coef_plot(b, error_bar = "se", order_by = "strength", to = "SB") + ggplot2::coord_flip()


# Averaging models within 2 CICc points of top model
avg <- average(P, avg_method = "full")
#Method full is because I'm using full averaging, where I include models that don't have the same paths
avg$coef

coef_plot(avg, error_bar = "se", order_by = "strength", to = "SB") + ggplot2::coord_flip()
coef_plot(avg, error_bar = "se", order_by = "strength", to = "BL") + ggplot2::coord_flip()
coef_plot(avg, error_bar = "se", order_by = "strength", to = "HP") + ggplot2::coord_flip()

coef_plot(avg)
plot(avg, curvature = 0.2)

P$d_sep$D


#############################################################################################
#############################################################################################

# Larger datasets looking at interactions

# Social, Colour, Body size (N = 229)
rm(list=ls())
library(phylopath)
library(ape)
require(geiger)
library(phytools)
setwd('C:\\Users\\cm15245\\OneDrive - University of Bristol\\Documents\\(0) PhD\\Comparative stuff\\Data\\(0) NEW datasets')

phy<-ape::read.nexus(file = 'SB_CP_BSTree.nex')

library(openxlsx)
excel <- read.xlsx(xlsxFile = "SB_CP_BS.xlsx")
head(excel)
excel$SB<-as.factor(excel$SB)
excel$CP<-as.factor(excel$CP)

SB <- excel$SB
names(SB) <- excel$phylo
CP<-excel$CP
names(CP)<-excel$phylo
BL<-excel$BS
names(BL)<-excel$phylo

data<-data.frame(SB,CP,BL)

str(data)

m5 <- define_model_set(
  null = c(),
  B = c(BL~CP, SB~BL),
  C = c(BL~CP, SB~CP),
  D = c(CP~BL, SB~CP),
  E = c(SB~CP, BL~SB),
  G = c(SB~CP+BL),
  H = c(SB~CP))  

P5<-phylo_path(model_set = m5, data = data, tree = phy)
P5
s5 <- summary(P5)
s5
plot(s5)

plot(m5$C)
plot(m5$D)

b <- best(P5) 
plot(b)
b_ci <- best(P5, boot = 500)
b_ci
plot(b_ci)

coef_plot(b, error_bar = "se", order_by = "strength", to = "SB") + ggplot2::coord_flip()
# Shows that colour pattern has a large effect on social behaviour in the best supported model

# Averaging models within 2 CICc points of top model
avg <- average(P5, avg_method = "full")
avg$coef

coef_plot(avg, error_bar = "se", order_by = "strength", to = "BS") + ggplot2::coord_flip()
coef_plot(avg, error_bar = "se", order_by = "strength", to = "CP") + ggplot2::coord_flip()

plot(avg, curvature = 0.1)

#------------------------------
# Social, colour and host plant (N = 217)

rm(list=ls())
library(phylopath)
library(ape)
require(geiger)
library(phytools)
setwd('C:\\Users\\cm15245\\OneDrive - University of Bristol\\Documents\\(0) PhD\\Comparative stuff\\Data\\(0) NEW datasets')

phy<-ape::read.nexus(file = 'CP_HPTree.nex')

library(openxlsx)
excel <- read.xlsx(xlsxFile = "SB_CP_HP.xlsx")
head(excel)
excel$SB<-as.factor(excel$SB)
excel$CP<-as.factor(excel$CP)
excel$HP<-as.factor(excel$HP)

SB <- excel$SB
names(SB) <- excel$phylo
CP <- excel$CP
names(CP) <- excel$phylo
HP<-excel$HP
names(HP)<-excel$phylo

data<-data.frame(SB,CP,HP)
str(data)

m2 <- define_model_set(
  null = c(),
  A = c(SB~CP, HP~CP),
  B = c(SB~CP, CP~HP),
  C = c(SB~CP+HP),
  D = c(SB~CP,HP~SB),
  E = c(HP~CP,SB~HP),
  F = c(SB~CP)) 


P2<-phylo_path(model_set = m2, data = data, tree = phy)
P2
s2 <- summary(P2)
s2
plot(s2)

plot(m2$A)
plot(m2$B)
plot(m2$D)
plot(m2$C)

b <- best(P2) 
plot(b)
b_ci <- best(P2, boot = 500)
b_ci
plot(b_ci)

coef_plot(b, error_bar = "se", order_by = "strength", to = "SB") + ggplot2::coord_flip()

# Averaging models within 2 CICc points of top model
avg <- average(P2, avg_method = "full")
avg$coef

coef_plot(avg, error_bar = "se", order_by = "strength", to = "HP") + ggplot2::coord_flip()

coef_plot(avg)
plot(avg, curvature = 0.4)
```
