## Supplemental R scripts for "Warning coloration, body size and the evolution of gregarious behavior in butterfly larvae": Transition_and_Coevolution.R

```
# Larval body size correlation analysis

rm(list=ls())
library(ape)
library(geiger)
library(phytools)
library(nlme)
library(openxlsx)
library(caper)
library(ggplot2)

setwd('C:\\Users\\cm15245\\OneDrive - University of Bristol\\Documents\\(0) PhD\\Comparative stuff\\Data\\(0) NEW datasets')
tree<-ape::read.nexus(file = 'LBS_WSTree.nex')
data <- read.xlsx(xlsxFile = "SB_WS_LBS.xlsx")
str(data)
data$SB<- as.factor(data$SB)

sizes <- comparative.data(phy=tree, data= data, names.col=phylo, vcv=TRUE, vcv.dim=3, na.omit=T)
#caper needs us to combine the table and tree into a new object to have the information together
sizes


PGLSmod <- pgls(WS ~ LBS, data=sizes, lambda ='ML', bounds=list(lambda=c(0,1)))
summary(PGLSmod)

# Plot the data
plot(PGLSmod)
coef(PGLSmod)

plot(data$WS ~ data$LBS)

abline(a = coef(PGLSmod)[1], b = coef(PGLSmod)[2])


# Plot real larval body size distribution
ggplot(data, aes(x=LBS)) + geom_histogram(binwidth = 5,color="black", fill="white")+ 
  scale_x_continuous(breaks=c(10,20,30,40,50,60,70,80,90,100,110,120,130,140), limits=c(0,140),
                     name = 'Final instar body length (mm)')+ scale_y_continuous(name="Frequency", limits=c(0, 20))+
  theme(panel.grid.major = element_blank(), panel.grid.minor = element_blank())+
  theme(panel.border = element_blank())+
  theme(axis.line = element_line(colour = "black"),panel.background = element_blank(),
        axis.title=element_text(size=15))


###############################################################################
###############################################################################

# Trait evolution analyses

# Transition rates using individual (larger) datasets

# Social behaviour
rm(list=ls())
library(ape)
require(geiger)
library(phytools)
library(openxlsx)
library(phylopath)
library(ggplot2)

setwd('C:\\Users\\cm15245\\OneDrive - University of Bristol\\Documents\\(0) PhD\\Comparative stuff\\Data\\(0) NEW datasets')
phylo<-ape::read.nexus(file = 'SB_onlyTree.nex')
excel <- read.xlsx(xlsxFile = "SB_only.xlsx")
excel$SB <- as.factor(excel$SB)
str(excel)
SB <- excel$SB
names(SB) <- excel$phylo
SB
str(SB)


SBAER<-fitDiscrete(phylo, SB,model = 'ER')
SBAER #AIC = 357.915168, log-likelihood = -177.957584
SBER1<-fitDiscrete(phylo, SB,transform = 'lambda', model = 'ER')
SBER1 #AIC = 358.084480, log-likelihood = -177.042240, lambda = 0.947451

lrstat<-2*(-177.042240-(-177.957584))
lrstat
pchisq(1.830688,1,lower.tail=F)
#0.1760463
# Lambda is not significant

# Then test for which model of evolution is best fitting
SBER<-fitDiscrete(phylo, SB, model = 'ER')
SBER #AIC = 357.915168, log-likelihood = -177.957584
SBARD<-fitDiscrete(phylo, SB,model = 'ARD')
SBARD #AIC = 353.960378, log-likelihood = -174.980189

lrstat<-2*(-174.980189-(-177.957584))
lrstat
pchisq(5.95479,1,lower.tail=F)
#0.01467735
# ARD model is best fitting. Where changes from greg to solitary happen faster than reverse
#     0          1
#0 -0.005863854  0.005863854
#1  0.019722989 -0.019722989

#Ancestral state
SBace<-make.simmap(phylo,SB,model="ARD",nsim=10000,pi="estimated",method='ML')
summary(SBace)
summary(SBace)$ace
#0     1
#0.8987 0.1013
countSimmap(SBace)

save(SBace, file = "SBace.RData")
load(file = "SBace.RData")

acenodes<- summary(SBace)$ace
par(mar=c(0,0,0,0))
plot(phylo, type="fan",label.offset = 4, cex = 0.4)
co <- c("purple","Tomato") 
tiplabels(pch = 16, col = co[as.numeric(SB)], cex = 0.4, adj = 1,offset = 3)
nodelabels(pie=acenodes,piecol = co[1:max(as.numeric(SB))], cex=0.2)
nodelabels(cex=0.2)
cladelabels(tree=NULL, text, node=5, offset=NULL, wing.length=NULL, cex=1)


#---------------------------
# Colour pattern
rm(list=ls())
library(ape)
require(geiger)
library(phytools)
library(openxlsx)
library(phylopath)
setwd('C:\\Users\\cm15245\\OneDrive - University of Bristol\\Documents\\(0) PhD\\Comparative stuff\\Data\\(0) NEW datasets')
phylo<-ape::read.nexus(file = 'SB_CPTree.nex')
excel <- read.xlsx(xlsxFile = "CP_only.xlsx")
excel$CP <- as.factor(excel$CP)
str(excel)
CP <- excel$CP
names(CP) <- excel$phylo

CPER<- fitDiscrete(phylo, CP, model = "ER")
CPER #AIC = 274.029900, log-likelihood = -136.014950
CPER1<- fitDiscrete(phylo, CP, model = "ER", transform = 'lambda')
CPER1 #AIC = 275.387281, log-likelihood = -135.693640, lambda = 0.961807
lrstat<-2*(-135.693640-(-136.014950))
lrstat
pchisq(0.64262,1,lower.tail=F)
#0.4227636 - Lambda is not significant

CPER<- fitDiscrete(phylo, CP, model = "ER")
CPER #AIC = 274.029900, log-likelihood = -136.014950
CPARD<- fitDiscrete(phylo, CP, model = "ARD")
CPARD
# AIC = 264.789069, log-likelihood = -130.394534 

lrstat<-2*(-130.394534-(-136.014950))
lrstat
pchisq(11.24083,1,lower.tail=F)
#0.0008001737 - ARD model is better fitting
# Apos to cryp is faster than reverse
#     0           1
#0 -0.002508624  0.002508624
#1  0.015506319 -0.015506319

#Ancestral state
Colace<-make.simmap(phylo,CP,model="ARD",nsim=10000,pi="estimated", method='ML')
save(Colace, file = "Colace.RData")
load(file = "Colace.RData")

summary(Colace)$ace
#0    1
#0.9472 0.0528
summary(Colace)

acenodes1<- summary(Colace)$ace
png('CP_onlyTree.png', width = 465, height = 465, units = 'mm', res = 300)
par(mar=c(0,0,0,0))
plot(phylo, type="fan",label.offset = 6, cex = 1)
co <- c("darkgrey","darkorange1") 
tiplabels(pch = 16, col = co[as.numeric(CP)], cex = 1.3, adj = 1,offset = 4.5)
nodelabels(pie=acenodes1,piecol = co[1:max(as.numeric(CP))], cex=0.2)
dev.off()

#---- SB and CP tree
rm(list=ls())
library(ape)
require(geiger)
library(phytools)
library(openxlsx)
library(phylopath)
setwd('C:\\Users\\cm15245\\OneDrive - University of Bristol\\Documents\\(0) PhD\\Comparative stuff\\Data\\(0) NEW datasets')
phylo<-ape::read.nexus(file = 'SB_CPTree.nex')
phylo
excel <- read.xlsx(xlsxFile = "SB_CP.xlsx") 
excel$SB<- as.factor(excel$SB)
SB <- excel$SB
names(SB) <- excel$phylo

SB_CPace<-make.simmap(phylo,SB,model="ARD",nsim=10000,pi="estimated", method='ML')
save(SB_CPace, file = "SB_CPace.RData")
load(file = "SB_CPace.RData")

summary(SB_CPace)$ace
#0    1
#0.9631 0.0369
summary(SB_CPace)

acenodes2<- summary(SB_CPace)$ace
png('SB_CPTree.png', width = 465, height = 465, units = 'mm', res = 300)
par(mar=c(0,0,0,0))
plot(phylo, type="fan",label.offset = 4, cex = 1)
co <- c("darkgrey","dodgerblue1") 
tiplabels(pch = 16, col = co[as.numeric(SB)], cex = 1.3, adj = 1,offset = 1.5)
nodelabels(pie=acenodes2,piecol = co[1:max(as.numeric(SB))], cex=0.3)
dev.off()

#---------------------------
# Structural defence
rm(list=ls())
library(ape)
require(geiger)
library(phytools)
library(openxlsx)
library(phylopath)
setwd('C:\\Users\\cm15245\\OneDrive - University of Bristol\\Documents\\(0) PhD\\Comparative stuff\\Data\\(0) NEW datasets')
phylo<-ape::read.nexus(file = 'SB_SDTree.nex')
excel <- read.xlsx(xlsxFile = "SD_only.xlsx")
excel$SD <- as.factor(excel$SD)
str(excel)
SD <- excel$SD
names(SD) <- excel$phylo

SDER<- fitDiscrete(phylo, SD, model = "ER")
SDER #AIC = 170.850389, log-likelihood = -84.425195
SDER1<- fitDiscrete(phylo, SD, model = "ER", transform = 'lambda')
SDER1 #AIC = 172.850389, log-likelihood = -84.425195, lambda = 1.000000
lrstat<-2*(-84.425195-(-84.425195))
lrstat
pchisq(0,1,lower.tail=F)
#1 - Lambda is not significant

SDER<- fitDiscrete(phylo, SD, model = "ER")
SDER #AIC = 170.850389, log-likelihood = -84.425195
SDARD<- fitDiscrete(phylo, SD, model = "ARD")
SDARD #AIC = 172.835281, log-likelihood = -84.417641
lrstat<-2*(-84.417641-(-84.425195))
lrstat
pchisq(0.015108,1,lower.tail=F)
#0.9021747 - ER model is better fitting


#Ancestral state
SDace<-make.simmap(phylo,SD,model="ER",nsim=10000,pi="estimated",method='ML')
save(SDace, file = "SDace.RData")
load(file = "SDace.RData")

summary(SDace)$ace
# 0    1
#0.9904 0.0096
summary(SDace)

#---------------------------
# Host plant - Categorical
rm(list=ls())
library(ape)
require(geiger)
library(phytools)
library(openxlsx)
library(phylopath)
setwd('C:\\Users\\cm15245\\OneDrive - University of Bristol\\Documents\\(0) PhD\\Comparative stuff\\Data\\(0) NEW datasets')
phylo<-ape::read.nexus(file = 'SB_HPTree.nex')
excel <- read.xlsx(xlsxFile = "HP_Cat_only.xlsx")
str(excel)
HP <- excel$HP
names(HP) <- excel$phylo
HP
str(HP)

HPER<- fitDiscrete(phylo, HP, model = "ER")
HPER #AIC = 631.850150, log-likelihood = -314.925075
HPER1<- fitDiscrete(phylo, HP, model = "ER", transform = 'lambda')
HPER1 #AIC = 630.983011, log-likelihood = -313.491506, lambda = 0.936748
lrstat<-2*(-313.491506-(-314.925075))
lrstat
pchisq(2.867138,1,lower.tail=F)
#0.09040551 - Lambda is not significant

HPER<- fitDiscrete(phylo, HP, model = "ER")
HPER #AIC = 631.850150, log-likelihood = -314.925075
HPSYM<- fitDiscrete(phylo, HP, model = "SYM")
HPSYM #AIC = 616.700632, log-likelihood = -298.350316
HPARD<- fitDiscrete(phylo, HP, model = "ARD")
HPARD #AIC = 601.545828, log-likelihood = -280.772914

lrstat<-2*(-298.350316-(-314.925075)) # ER vs SYM model
lrstat
pchisq(33.14952,9,lower.tail=F)
#0.0001257784 - Significant, so can use more complex SYM model
lrstat<-2*(-280.772914-(-298.350316)) # SYM vs ARD model
lrstat
pchisq(35.1548,10,lower.tail=F)
#0.0001174926 - Significant, so use more complex ARD model
# Fastest transition denoted with asterisk

#             Forb/herb     Graminoid         Shrub        Tree          Vine
#Forb/herb -1.940245e-02  1.328575e-03  1.259818e-02*  0.005475699  7.624210e-17
#Graminoid  6.963263e-03* -1.362105e-02  1.640619e-06  0.006629079  2.706799e-05
#Shrub      5.593900e-07  6.779809e-07 -6.040550e-02  0.038499127*  2.190513e-02
#Tree       5.215897e-03  1.754226e-07  6.378928e-03* -0.011595001  2.894907e-11
#Vine       6.321877e-03  1.189818e-06  9.654569e-17  0.014830779* -2.115385e-02

#Ancestral state
HPace<-make.simmap(phylo,HP,model="ARD",nsim=10000,pi="estimated",method='ML')
save(HPace, file = "HPace.RData")
load(file = "HPace.RData")

summary(HPace)$ace
# Forb/herb* Graminoid  Shrub   Tree   Vine
#  0.6452    0.0003    0.0048   0.3415  0.0082
summary(HPace)
#------
# Host plant - Binary (0-Tree, 1-Not tree)
rm(list=ls())
library(ape)
require(geiger)
library(phytools)
library(openxlsx)
library(phylopath)
setwd('C:\\Users\\cm15245\\OneDrive - University of Bristol\\Documents\\(0) PhD\\Comparative stuff\\Data\\(0) NEW datasets')
phylo<-ape::read.nexus(file = 'SB_HPTree.nex')
excel <- read.xlsx(xlsxFile = "HP_Bin.xlsx")
excel$HP <- as.factor(excel$HP)
str(excel)
HP <- excel$HP
names(HP) <- excel$phylo


HPER<- fitDiscrete(phylo, HP, model = "ER")
HPER #AIC = 307.124238, log-likelihood = -152.562119
HPER1<- fitDiscrete(phylo, HP, model = "ER", transform = 'lambda')
HPER1 #AIC = 306.280094, log-likelihood = -151.140047, lambda = 0.913829
lrstat<-2*(-151.140047-(-152.562119))
lrstat
pchisq(2.844144,1,lower.tail=F)
#0.09170742 - Lambda is not significant

HPER<- fitDiscrete(phylo, HP, model = "ER")
HPER #AIC = 307.124238, log-likelihood = -152.562119
HPARD<- fitDiscrete(phylo, HP, model = "ARD")
HPARD #AIC = 308.804910, log-likelihood = -152.402455
lrstat<-2*(-152.402455-(-152.562119))
lrstat
pchisq(0.319328,1,lower.tail=F)
#0.5720118 - ER model is better fitting


#Ancestral state
HPace2<-make.simmap(phylo,HP,model="ER",nsim=10000,pi="estimated",method='ML')
save(HPace2, file = "HPace2.RData")
load(file = "HPace2.RData")

summary(HPace2)$ace # No difference
# 0    1
#0.4973 0.5027
summary(HPace2)


#----------------
# Body size
rm(list=ls())
library(ape)
library(geiger)
library(phytools)
library(openxlsx)
library(phylopath)
library(phylolm)
library(picante)
library(caper)
library(mmeln)

setwd('C:\\Users\\cm15245\\OneDrive - University of Bristol\\Documents\\(0) PhD\\Comparative stuff\\Data\\(0) NEW datasets')
phylo<-ape::read.nexus(file = 'SB_BSTree.nex')
phylo
excel <- read.xlsx(xlsxFile = "WS_only.xlsx")
str(excel)

BS<-excel$BS
names(BS)<-excel$phylo
BS

phylosig(phylo,BS,test=TRUE)
# K = 0.835216 
# p = 0.001
phylosig(phylo,BS,method="lambda",test=TRUE)
# Lambda = 0.936352
# p = 5.36487e-51
fitBM<-fitContinuous(phylo,BS)#169.819672
fitOU<-fitContinuous(phylo,BS,model="OU")#171.383417
fitEB<-fitContinuous(phylo,BS,model="EB")#169.819393
aic.vals<-setNames(c(fitBM$opt$aicc,fitOU$opt$aicc,fitEB$opt$aicc),
                   c("BM","OU","EB"))
aic.vals
# Best supported model of trait evolution is BM

lrstat<-2*(171.383417-(169.819672)) #BM vs OU
lrstat
pchisq(3.12749,1,lower.tail=F)
#0.07698218 BM better fitting
lrstat<-2*(169.819672-(169.819393)) #BM vs EB
lrstat
pchisq(0.000558,1,lower.tail=F)
#0.9811541 BM better fitting

#Ancestral state
WSace<- ace(BS, phylo, type = 'continuous', method = 'ML', CI = TRUE, model = 'BM')
WSace$ace
#1.722804
WSace
# lower    upper
#1.663782 1.781827


#############################################################################
#############################################################################

# Coevolution with larger, individual datasets
rm(list=ls())
library(ape)
library(geiger)
library(phytools)
library(openxlsx)
library(phylopath)
library(phylolm)
library(picante)
library(caper)

##Social behaviour VS Colour pattern
setwd('C:\\Users\\cm15245\\OneDrive - University of Bristol\\Documents\\(0) PhD\\Comparative stuff\\Data\\(0) NEW datasets')
phy<-ape::read.nexus(file = 'SB_CPTree.nex')
phy
excel <- read.xlsx(xlsxFile = "SB_CP.xlsx") 
str(excel)
excel$SB<- as.factor(excel$SB)
excel$CP<- as.factor(excel$CP)
SB <- excel$SB
names(SB) <- excel$phylo
CP<-excel$CP
names(CP)<-excel$phylo


SBVCP_pagel <- fitPagel(phy,SB,CP, model = 'ARD', dep.var = "x")
save(SBVCP_pagel, file = "SBVCP_pagel.RData")
SBVCP_pagel$independent.logL
#-247.3815
SBVCP_pagel$dependent.logL
#-235.9135
SBVCP_pagel$independent.AIC
#502.7631
SBVCP_pagel$dependent.AIC
#483.8271
plot(SBVCP_pagel)
#looking at the sides
#Solitary to greg is faster when aposematic
#Greg to solitary is faster when cryptic

lrstat<-2*(-235.9135-(-247.3815))
lrstat
pchisq(22.936,2,lower.tail=F)
# 1.04595e-05

SBVCP_pagel1 <- fitPagel(phy,SB,CP, model = 'ARD', dep.var = "y")
save(SBVCP_pagel1, file = "SBVCP_pagel1.RData")
load(file = "SBVCP_pagel1.RData")
SBVCP_pagel1$independent.logL
#-247.3815
SBVCP_pagel1$dependent.logL
#-236.218
SBVCP_pagel1$independent.AIC
#502.7631
SBVCP_pagel1$dependent.AIC
#484.4361
plot(SBVCP_pagel1)
#Looking at top and bottom
#Cryp to apos is faster when gregarious
#Apos to cryp is faster when solitary

lrstat<-2*(-236.218-(-247.3815))
lrstat
pchisq(22.327,2,lower.tail=F)
# 1.418252e-05

SBVCP_pagel2 <- fitPagel(phy,SB,CP, model = 'ARD', dep.var = "xy")
save(SBVCP_pagel2, file = "SBVCP_pagel2.RData")
SBVCP_pagel2$independent.logL
#-247.3815
SBVCP_pagel2$dependent.logL
#-233.4336
SBVCP_pagel2$independent.AIC
#502.7631
SBVCP_pagel2$dependent.AIC
#482.8672
plot(SBVCP_pagel2)


#-----------------------

##Social behaviour VS Structural defence
rm(list=ls())
library(ape)
setwd('C:\\Users\\cm15245\\OneDrive - University of Bristol\\Documents\\(0) PhD\\Comparative stuff\\Data\\(0) NEW datasets')
phy<-ape::read.nexus(file = 'SB_SDTree.nex')
phy
excel <- read.xlsx(xlsxFile = "SB_SD.xlsx") 
excel$SB<- as.factor(excel$SB)
excel$SD<- as.factor(excel$SD)
SB <- excel$SB
names(SB) <- excel$phylo
str(SB)
SD<-excel$SD
names(SD)<-excel$phylo

SBVSD_pagel <- fitPagel(phy,SB,SD, model = 'ARD',dep.var = "x")
save(SBVSD_pagel, file = "SBVSD_pagel.RData")
load(file = "SBVSD_pagel.RData")
SBVSD_pagel$independent.logL
#-182.494
SBVSD_pagel$dependent.logL
#-179.0519
SBVSD_pagel$independent.AIC
#372.9879
SBVSD_pagel$dependent.AIC
#370.1038
plot(SBVSD_pagel)
#Dependent model, sides
# Sol to greg faster with defence
# Greg to sol faster without defence

lrstat<-2*(-179.0519-(-182.494))
lrstat
pchisq(6.8842,2,lower.tail=F)
# 0.03199742

SBVSD_pagel1 <- fitPagel(phy,SB,SD, model = 'ARD',dep.var = "y")
save(SBVSD_pagel1, file = "SBVSD_pagel1.RData")
SBVSD_pagel1$independent.logL
#-182.494
SBVSD_pagel1$dependent.logL
#-176.6586
SBVSD_pagel1$independent.AIC
#372.9879
SBVSD_pagel1$dependent.AIC
#365.3173
plot(SBVSD_pagel1)
#Dependent model. Looking at top and bottom
#No Defence to Defence faster when gregarious
#Defence to No Defence is faster when solitary

lrstat<-2*(-176.6586-(-182.494))
lrstat
pchisq(11.6708,2,lower.tail=F)
# 0.002922254

#-----------------------

# Social behaviour Vs Host plant
rm(list=ls())
library(ape)
setwd('C:\\Users\\cm15245\\OneDrive - University of Bristol\\Documents\\(0) PhD\\Comparative stuff\\Data\\(0) NEW datasets')
phy<-ape::read.nexus(file = 'SB_HPTree.nex')
phy
excel <- read.xlsx(xlsxFile = "SB_HP.xlsx")  
head(excel)
excel$SB<- as.factor(excel$SB)
excel$HP<- as.factor(excel$HP)
SB <- excel$SB
names(SB) <- excel$phylo
str(SB)
HP<-excel$HP
names(HP)<-excel$phylo

SBVHP_pagel <- fitPagel(phy,SB,HP, model = 'ARD', dep.var = "x")
save(SBVHP_pagel, file = "SBVHP_pagel.RData")
SBVHP_pagel$independent.logL
#-280.6393
SBVHP_pagel$dependent.logL
#-279.6083
SBVHP_pagel$independent.AIC
#569.2786
SBVHP_pagel$dependent.AIC
#571.2165
plot(SBVHP_pagel)
#Independent model

lrstat<-2*(-279.6083-(-280.6393))
lrstat
pchisq(2.062,2,lower.tail=F)
# 0.3566501

SBVHP_pagel1 <- fitPagel(phy,SB,HP, model = 'ARD', dep.var = "y")
save(SBVHP_pagel1, file = "SBVHP_pagel1.RData")
SBVHP_pagel1$independent.logL
#-280.6393
SBVHP_pagel1$dependent.logL
#-280.0755
SBVHP_pagel1$independent.AIC
#569.2786
SBVHP_pagel1$dependent.AIC
#572.151
plot(SBVHP_pagel1)
#Independent model

lrstat<-2*(-280.0755-(-280.6393))
lrstat
pchisq(1.1276,2,lower.tail=F)
# 0.5690426


#-----------------------------------------------------------------------

##Colour pattern VS Structural defence
rm(list=ls())
library(ape)
setwd('C:\\Users\\cm15245\\OneDrive - University of Bristol\\Documents\\(0) PhD\\Comparative stuff\\Data\\(0) NEW datasets')
phy<-ape::read.nexus(file = 'CP_SDTree.nex')
phy
excel <- read.xlsx(xlsxFile = "CP_SD.xlsx")
excel$CP<- as.factor(excel$CP)
excel$SD<- as.factor(excel$SD)
CP <- excel$CP
names(CP) <- excel$phylo
SD<-excel$SD
names(SD)<-excel$phylo

CPVSD_pagel <- fitPagel(phy,CP,SD, model='ARD', dep.var = "x")
save(CPVSD_pagel, file = "CPVSD_pagel.RData")
CPVSD_pagel$independent.logL
#-191.5799
CPVSD_pagel$dependent.logL
#-183.5791
CPVSD_pagel$independent.AIC
#391.1598
CPVSD_pagel$dependent.AIC
#379.1582
plot(CPVSD_pagel)
#Dependent model, sides
#Cryp to apos faster with defence
#Apos to cryp faster without defence

lrstat<-2*(-183.5791-(-191.5799))
lrstat
pchisq(16.0016,2,lower.tail=F)
# 0.0003351944

CPVSD_pagel1 <- fitPagel(phy,CP,SD, model='ARD', dep.var = "y")
save(CPVSD_pagel1, file = "CPVSD_pagel1.RData")
CPVSD_pagel1$independent.logL
#-191.5799
CPVSD_pagel1$dependent.logL
#-179.7706
CPVSD_pagel1$independent.AIC
#391.1598
CPVSD_pagel1$dependent.AIC
#371.5412
plot(CPVSD_pagel1)
#Dependent model. Looking at top and bottom
#No Defence to Defence faster when apos
#Defence to No Defence is faster when cryptic

lrstat<-2*(-179.7706-(-191.5799))
lrstat
pchisq(23.6186,2,lower.tail=F)
# 7.435089e-06


#-----------------------

##Colour pattern VS Host plant
rm(list=ls())
library(ape)
setwd('C:\\Users\\cm15245\\OneDrive - University of Bristol\\Documents\\(0) PhD\\Comparative stuff\\Data\\(0) NEW datasets')
phy<-ape::read.nexus(file = 'CP_HPTree.nex')
phy
excel <- read.xlsx(xlsxFile = "CP_HP_Bin.xlsx")
excel
excel$CP<- as.factor(excel$CP)
excel$HP<- as.factor(excel$HP)
CP<-excel$CP
names(CP)<-excel$phylo
HP<-excel$HP
names(HP)<-excel$phylo

CPVHP_pagel <- fitPagel(phy,CP,HP, model='ARD', dep.var = "x")
save(CPVHP_pagel, file = "CPVHP_pagel.RData")
CPVHP_pagel$independent.logL
#-235.1899
CPVHP_pagel$dependent.logL
#-224.6753
CPVHP_pagel$independent.AIC
#478.3799
CPVHP_pagel$dependent.AIC
#461.3506
plot(CPVHP_pagel)
#Dependent model, sides
# Cryp to apos faster not on trees
# Apos to cryp also faster not on trees

lrstat<-2*(-224.6753-(-235.1899))
lrstat
pchisq(21.0292,2,lower.tail=F)
# 2.713734e-05


CPVHP_pagel1 <- fitPagel(phy,CP,HP, model='ARD', dep.var = "y")
save(CPVHP_pagel1, file = "CPVHP_pagel1.RData")
CPVHP_pagel1$independent.logL
#-235.1899
CPVHP_pagel1$dependent.logL
#-233.2983
CPVHP_pagel1$independent.AIC
#478.3799
CPVHP_pagel1$dependent.AIC
#478.5967
plot(CPVHP_pagel1)
#Independent model

lrstat<-2*(-233.2983-(-235.1899))
lrstat
pchisq(3.7832,2,lower.tail=F)
# 0.1508303

#-----------------------------------------------------------

##Structural defence VS Host plant
rm(list=ls())
library(ape)
setwd('C:\\Users\\cm15245\\OneDrive - University of Bristol\\Documents\\(0) PhD\\Comparative stuff\\Data\\(0) NEW datasets')
phy<-ape::read.nexus(file = 'SD_HPTree.nex')
phy
excel <- read.xlsx(xlsxFile = "SD_HP.xlsx",sheet = 1)
excel
excel$SD<- as.factor(excel$SD)
excel$HP<- as.factor(excel$HP)
SD<-excel$SD
names(SD)<-excel$phylo
HP<-excel$HP
names(HP)<-excel$phylo

SDVHP_pagel <- fitPagel(phy,SD,HP, model='ARD', dep.var = "x")
save(SDVHP_pagel, file = "SDVHP_pagel.RData")
SDVHP_pagel$independent.logL
#-185.4882
SDVHP_pagel$dependent.logL
#-185.0392
SDVHP_pagel$independent.AIC
#378.9765
SDVHP_pagel$dependent.AIC
#382.0785
plot(SDVHP_pagel)
# independent model

lrstat<-2*(-185.0392-(-185.4882))
lrstat
pchisq(0.898,2,lower.tail=F)
# 0.6382661

SDVHP_pagel1 <- fitPagel(phy,SD,HP, model='ARD', dep.var = "y")
save(SDVHP_pagel1, file = "SDVHP_pagel1.RData")
SDVHP_pagel1$independent.logL
#-185.4882
SDVHP_pagel1$dependent.logL
#-185.0749
SDVHP_pagel1$independent.AIC
#378.9765
SDVHP_pagel1$dependent.AIC
#382.1497
plot(SDVHP_pagel1)
# independent model

lrstat<-2*(-185.0749-(-185.4882))
lrstat
pchisq(0.8266,2,lower.tail=F)
# 0.6614638
```
